# The Interaction of NF-κB Transcription Factor with Centromeric Chromatin

**DOI:** 10.1101/2024.02.13.580208

**Authors:** Shaun Filliaux, Chloe Bertelsen, Hannah Baughman, Elizabeth Komives, Yuri L. Lyubchenko

## Abstract

Centromeric chromatin is a subset of chromatin structure and governs chromosome segregation. The centromere is composed of both CENP-A nucleosomes (CENP-A_nuc_) and H3 nucleosomes (H3_nuc_) and is enriched with alpha-satellite (α-sat) DNA repeats. These CENP-A_nuc_ have a different structure than H3_nuc_, decreasing the base pairs (bp) of wrapped DNA from 147 bp for H3_nuc_ to 121 bp for CENP-A_nuc_. All these factors can contribute to centromere function. We investigated the interaction of H3_nuc_ and CENP-A_nuc_ with NF-κB, a crucial transcription factor in regulating immune response and inflammation. We utilized Atomic Force Microscopy (AFM) to characterize complexes of both types of nucleosomes with NF-κB. We found that NF-κB unravels H3_nuc_, removing more than 20 bp of DNA, and that NF-κB binds to the nucleosomal core. Similar results were obtained for the truncated variant of NF-κB comprised only of the Rel Homology domain and missing the transcription activation domain (TAD), suggesting the RelA TAD is not critical in unraveling H3_nuc_. By contrast, NF-κB did not bind to or unravel CENP- A_nuc_. These findings with different affinities for two types of nucleosomes to NF-κB may have implications for understanding the mechanisms of gene expression in bulk and centromere chromatin.

## Introduction

The centromere is a specialized chromatin region located at the center of each eukaryotic chromosome and is responsible for ensuring proper segregation during mitosis and meiosis ^1^. It consists of a complex network of proteins, DNA sequences, and chromatin structures that interact to form a cohesive unit responsible for accurately segregating chromosomes performed by the kinetochores ^2,3^. Although bulk chromatin consists of one type of nucleosomes with octamers consisting of duplicates of H2A, H2B, H3, and H4 histones (nucleosome H3_nuc_), centromere chromatin consists of two types of nucleosomes CENP-A (CENP-A_nuc_) and canonical H3_nuc_ nucleosomes ^4^. The only difference in these two nucleosomes is that in the H3_nuc_, histones are replaced with variant CENP-A histones. The centromere of most higher eukaryotes is comprised of alpha satellite (α-sat) motifs of a 171 bp DNA sequence ^5–7^. It is a biologically relevant sequence that is tandemly repeated hundreds to thousands of times, comprising 0.2-5 Mb stretches depending on the chromosome ^8–10^.

H3_nuc_ nucleosomes are composed of 147 bp DNA wrapped around a protein core of histone proteins (H2A, H2B, H3, and H4) and compact the genome into a more manageable structure ^11–13^. This compact structure is essential in protecting the genome from damage and plays a critical role in gene regulation ^14–17^.

In centromeric CENP-A octameric nucleosomes, H3 histones are replaced with CENP-A histones, an H3 homolog ^1,10,18–20^. These two homologs share a 50% homology in the C-terminal histone fold domain but vary drastically in the N-terminal tail in both size and sequence ^21,22^. These and other structural differences between CENP-A and H3 result in an unfixed 13 bp at both entry/exit of the CENP-A nucleosome, so centromeric CENP-A_nuc_ octameric nucleosomes wrap ∼ 20 bp less DNA than bulk H3 nucleosome ^23,24^.

Studies show that transcription does occur in the centromere, yielding different products depending on the number of repeats of a-sat ^25,26^. The transcription occurring in the centromere yields lncRNAs, which functionally load both CENP-A and CENP-C ^20^. However, there is a 200-300 fold difference between bulk chromatin and centromere transcription ^10,20,27^.

The accessibility of transcription factors to the bulk vs. centromere chromatin may be one of the explanations, so to test this hypothesis, we investigated the interaction of NF-κB transcription factor with both types of nucleosomes. NF-κB is a transcription factor that recognizes κB sites in the DNA and is crucial in regulating the immune response and inflammation. The α-sat sequence contains many half κB sites, which are also known to bind NF-κB ^10,20,28,29^. We previously reported that NF-κB binds and unravels H3_nuc_ nucleosomes assembled with the Widom 601 DNA motif ^30^. Here, we tested how NF-κB interacts with nucleosomes assembled on the centromere-specific a-sat sequence. Both H3_nuc_ and CENP-A_nuc_ were assembled on the same DNA substrate, and the interaction of the nucleosomes with NF-κB was studied using AFM. Analysis of AFM data revealed that NF-κB unravels H3_nuc_ but does not appear to bind or unravel CENP-A_nuc_ nucleosomes.

## Experimental Methods

### DNA preparation

The DNA construct was prepared the same way as we have done previously.^30–32^ The alpha satellite-containing construct was made using PCR with a pUC57 plasmid vector from BioBasic (Markham, ON, CA). The DNA total sequence was 410 bp, with the alpha satellite sequence in the middle. The specific sequence used is: 5’- GATGTGCTGCAAGGCGATTAAGTTGGGTAACGCCAGGGTTTTCCCAGTCACGACGTTGTAAAACGACGGCCAGTGAATTCGAGCTCGGTACCTCGCGAATGCATCTAGATGACCATTGGATTGAACTAACAGAGCTGAACACTCCTTTAGATGGAGCAGATTCCAAACACACTTTCTGTAGAATCTGCAAGTGGATATTTGGACTTCTCTGAGGATTTCGTTGGAAACGGGATAAAATTCCCAGAACTACACGGAAGCATTCTCAGAAACTTCTTTGTGATGAAGGGCGAATTCGAATCGGATCCCGGGCCCGTCGACTGCAGAGGCCTGCATGCAAGCTTGGCGTAATCATGGTCATAGCTGTTTCCTGTGTGAAATTGTTATCCGCTCACAATTCCACACAACATACG -3’. After the PCR amplification of the DNA substrate, the DNA was concentrated and purified using the Gel Extraction Kit from Qiagen (Hilden, DE). Lastly, the DNA concentrations were calculated using a NanoDrop Spectrophotometer (ND-1000, Thermo Fischer).

### Preparation of proteins (NF-κB)

N-terminal hexahistidine murine p50_39–350_/RelA_19–321_ (hereafter referred to as NF-κB_RHD_) was expressed using a modified pET22b vector containing the genes for both polypeptides as described previously ^33^. The DNA for murine RelA residues 19-549 was synthesized and subcloned into a modified pET22b vector which already contained the gene for N-terminal hexahistidine-p50_39-350_ (hereafter referred to as NF-κBA_FL_). The DNA sequence of the RelA_TAD_ (RelA residues 340-549) was subcloned into pET28a vector with a C-terminal hexahistidine tag.

All vectors were transformed into E. coli BL-21 (DE3) cells and grown to an OD_600_ of 0.5-0.7 at 37°C in M9 minimal media with antibiotic selection. Cultures were cooled on ice for 20 minutes, then protein expression was initiated by the addition of 0.2 mM IPTG. Cultures were incubated at 18°C for 16 hours, then harvested by centrifugation. Pellets were stored at -80°C.

The NF-κB_RHD_, NF-κB_FL_, and RelA_TAD_ constructs were lysed by sonication and purified by Ni^2+^-NTA chromatography as described previously for NF-κB_RHD_ ^30^. Following overnight dialysis, protein was aliquoted and stored at -80°C. Prior to experiments, aliquots were thawed and further purified. NF-κB_RHD_ and NF-κB_FL_ were purified by cation exchange chromatography (MonoS; GE healthcare) to remove bound nucleic acids, as described previously ^30^. Protein was further purified by size-exclusion chromatography using a Superdex 200 column (GE healthcare) in SEC buffer (25 mM Tris, 150 mM NaCl, 0.5 mM EDTA, 1 mM DTT, adjusted to pH 7.5 at room temperature). Care was taken to separate NF-κB_FL_ from a breakdown product that eluted at the same volume as NF-κB_RHD_. RelA_TAD_ was purified by size-exclusion chromatography using a Superdex 75 column, followed by a Superdex 200 column (GE healthcare) in the same buffer.

All purification chromatography steps were conducted in a 4°C cold room. Purity of all proteins was assessed by SDS-PAGE. The protein concentration was determined by absorption at 280 nm using a NanoDrop spectrophotometer. Purified protein was stored at 4°C and all experiments were conducted within 72 hours of purification by size exclusion chromatography.

### NF-κB reaction

The addition of NF-κB to DNA or nucleosome-containing samples was completed in the same manner as previously.^30^ The NF-κB was diluted to 300 nM for nucleosome experiments and 600 nM for DNA experiments in NF-κB buffer (25 mM Tris pH 7.5, 150 mM NaCl, 0.5 mM EDTA, 1 mM DTT). The DNA experiments incubated NF-κB at a 2:1 ratio with the DNA for 10 minutes at room temperature. The nucleosome experiments incubated NF-κB at a 1:1 ratio, resulting in 150 nM for both nucleosomes and NF-κB, and incubated for 10 minutes at room temperature.

### Nucleosome assembly

The nucleosome assembly utilizes a previously used method of dialyzing from a high salt concentration (2 M) to a low salt concentration of 2.5 mM. ^24^ Our nucleosome assembly starts in an initial buffer (10 mM Tris pH 7.5, 2M NaCl, 1 mM EDTA, 2 mM DTT), where we incubate the nucleosome octamers purchased from The Histone source (Fort Collins, CO) in a dialysis tube for one hour to allow the glycerol concentration to decrease. After the initial 1 hour incubation at 4 C, we start pumping the secondary buffer (10 mM Tris pH 7.5, 2.5 mM NaCl, 1 mM EDTA, 2 mM DTT) into the initial buffer using a peristaltic pump, which simultaneously pumps the initial buffer out, maintaining a consistent volume. The changing of NaCl concentration occurs for 24 hours at 4C. After the 24 hours are completed, another 1-hour incubation occurs in the secondary buffer to ensure the final concentration of NaCl is 2.5 mM. The CENP-A_nuc_ requires an additional step of adding in 1:2 tetramer:dimer molar concentrations for a proper octamer assembly. The CENP-A/H4 tetramer and H2A/H2B dimer are ordered from EpiCypher (Durham, NC)

### AFM imaging

The preparation of the samples was completed as performed previously by our lab. ^30,31^. The nucleosomes are stored at 300 nM, and the imaging of nucleosomes is completed at 2 nM. A dilution in our imaging buffer (4 mM MgCl_2_ and 10 mM HEPES) down to 2 nM is achieved for the control samples before the deposition. For samples that contain NF-κB, a mixture of the stock nucleosome sample and the NF-κB is done at the highest possible concentration. The 300 nM nucleosome stock is mixed with a 1:1 or 1:2 DNA: NF-κB ratio and incubated at room temperature for 10 minutes before diluting and preparing for deposition. AFM samples are deposited on functionalized APS mica, incubated for 2 minutes, washed with DI water, and gently dried under argon flow. Samples were stored in a vacuum before being imaged on a Multimode AFM/Nanoscope IIId system using TESPA probes (Bruker Nano Inc., Camarillo, CA).

### Data analysis

Data analysis was completed using previously successful techniques in our lab. ^30^ The contour length measurements are conducted using Femtoscan (Advance Technologies Center, Moscow, Russia). The measurements start at the DNA’s end and end in the middle of the protein (nucleosome or NF-κB). Then, the other DNA flank is measured. 5 nm is subtracted from each DNA flank to account for the length contributed by the histone core. The measurements are measured in nm and converted into bp through a conversion factor calculated by measuring naked DNA on each image. We take the full-length measurement of naked DNA and divide it by the known bp of the DNA to get a conversion factor typically around 0.35. Once DNA measurements are completed, Origin is used to fit the data into bins and the visual representation of histograms. Origin was used to calculate the mean gaussian distributions of the histograms. The mapping of the nucleosomes utilizes the two DNA flank measurements, and the short lengths were used to create bins. The nucleosome binding locations were created using Microsoft Excel from the bin information provided by Origin. The height measurements were calculated in Femtoscan using grain analysis and individually selecting each nucleosome. A height is calculated through the utilization of multiple cross-sections.

## Results

### NF-κB binding to the alpha satellite DNA substrate

In these studies, we used a DNA construct containing an alpha satellite (α-sat) sequence, 171 bp long, present exclusively in the centromere part of the chromosome flanked with the DNA segments that are not specific for nucleosome binding. Schematics of the DNA are shown in Figure 1A. α-sat segment was placed in the middle of the construct, indicated with a gray bar below. Green and orange bars indicate the positions of the NF-κB half recognition sequences. Both the central α-sat segment and flanks contain NF-κB binding sites.

**Figure 1.**
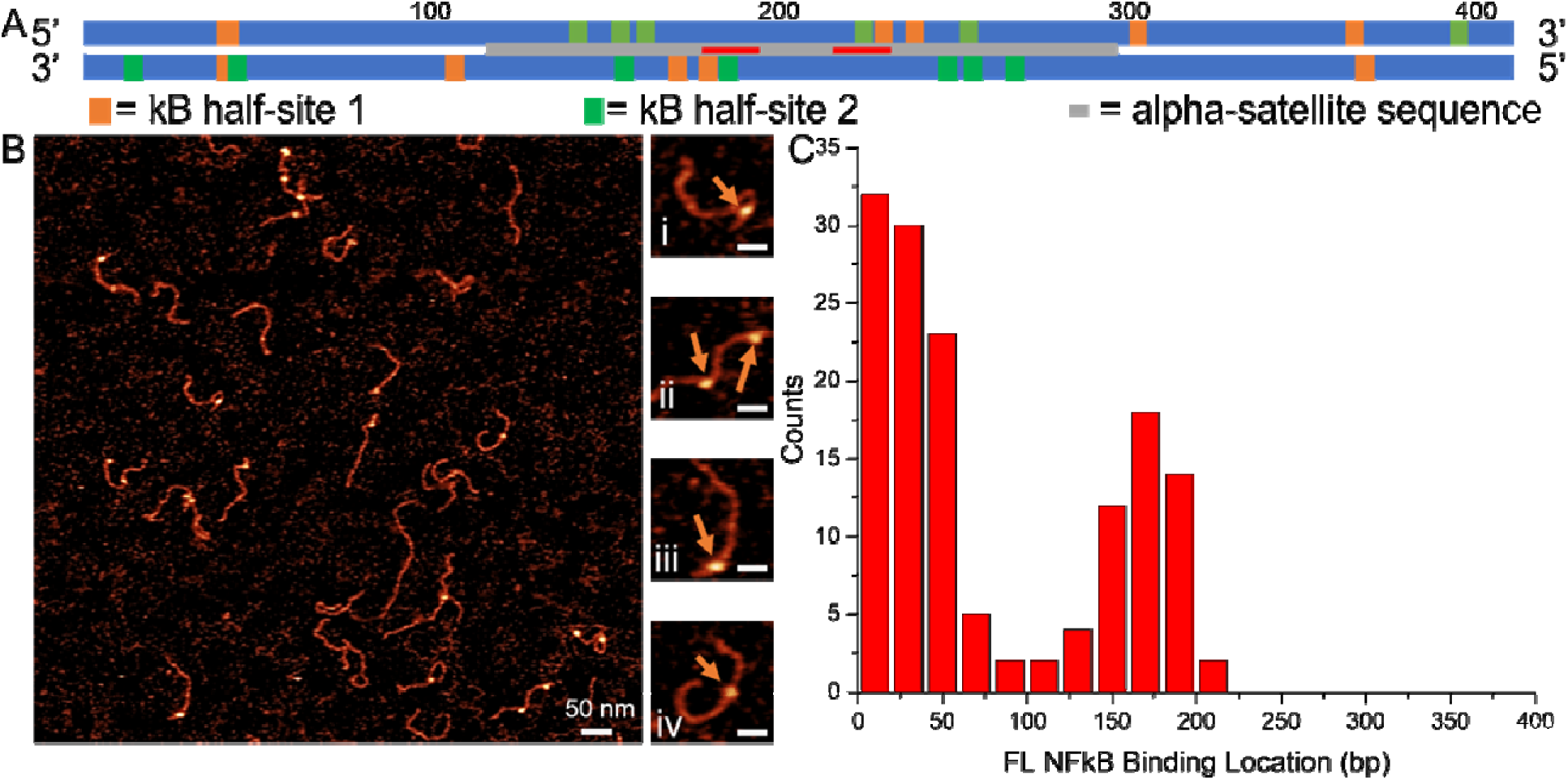
DNA Construct and AFM image with NF-κB_FL_ results. DNA construct (A) containing the alpha-satellite sequence in the middle of the sequence (grey bar), to analyze if there was any preferential binding at half sites, they are displayed in the “forward” 5’ -> 3’ direction and again in “reverse” 3’ ->5’, and the half-sites were marked (green and orange). The **NF-**κB half-sites 1 (GGGRN) were labeled with orange boxes, and the κB half-sites 2 (YYYCC) were labeled with green boxes. The results indicate a large cluster of half-sites near the middle of the DNA strand. The CENP-B box was marked in red. AFM images of NF-κB_FL_ at a 1 to 2 ratio(B) binding on the DNA substrate. Snapshots shown to the right of large AFM images (i, iii, and iv) show a single NF-κB_FL_ bound to the DNA, and image (ii) shows two NF-κB_FL_ bound to the DNA. The orange arrows indicate an NF-κB_FL_ bound to the DNA. The large AFM image i a 1 x 1 mm scan size with a 50 nm scale bar. The snapshots are 100 x 100 nm scan area and 25 nm scale bars. The binding location results for a single NF-κB_FL_ can be seen in the histogram (C), indicating a preference for terminal and a second peak around ∼170 bp.

The DNA was complexed with NF-κB_FL_ using a 1:1 protein-to-DNA molar ratio and imaged with AFM. AFM images are shown in Figure 1B. Protein bound to DNA appears as a globular feature on the DNA filament. Similar to our previous study ^30^, NF-κB does not alter the length of the DNA, suggesting that there is no wrapping DNA around the protein. A few zoomed images are displayed to the right (frames i-iv), and the protein position is indicated with arrows in these images. Complexes of the DNA with one or two NF-κB_FL_ molecules are seen, and both types of complexes are shown in selected frames. Protein can appear close to the end of the DNA (frames i and iii) or inside the DNA (frame iv). A similar arrangement was observed for two protein molecules bound to DNA (frames ii). Locations of the protein were mapped, and the results are shown as a histogram in Figure 1C. Two peaks correspond to the NF-κB_FL_ binding to the DNA ends (0-50 bp) and the central location (∼ 170 bp). The mapping results correlate with NF-κB binding sites shown in Figure 1A. Note that the left and right ends of the DNA cannot be distinguished in the AFM images, so the end-bound peak corresponds to complexes of NF-κB with left-right binding sites on the DNA construct. Similarly, the peak at 170 bp corresponds to NF-κB bound to half-κB sites (green or orange bars) on both DNA strands between 100 and 200 bp and to 200-300 segments of the DNA. The binding affinity for NF-κB between the peaks is low, which is in line with the lack of NF-κB sites on both DNA strands in the construct. These results agree with other papers demonstrating the need for a half κB site for binding ^34,35^.

Similar experiments were performed with the NF-κB_RHD_ variant in which the 228 amino acids of the TAD region were deleted. Regardless of the deletion, NF-κB_RHD_ demonstrates sequence- specific affinity very similar to the one for the full-length NF-κB heterodimer (Supplementary Fig. S1). These findings suggest that the C-terminal RelA TAD is not critical for the interaction of the NF-κB heterodimer with the DNA.

### NF-κB interaction with canonical nucleosomes H3_nuc_

Canonical nucleosomes H3_nuc_ were assembled on the DNA substrate described above by the self-assembly process described in the methods section using an octameric histone core containing H2A, H2B, H3, and H4. The AFM images of nucleosomes are shown in Figure 2A, with a few selected frames to the right of the large scan. Nucleosomes are indicated with blue arrows. In addition to terminal locations in frames (ii and iv), nucleosomes occupy positions near the middle of the sequence (frames i and iii). The various positions of the nucleosomes located in the middle of the α-sat segment demonstrate that the α-sat segment is not a nucleosome-specific sequence. This conclusion aligns with our previous publication in which a similar DNA substrate was used. ^31^ We also measured another parameter of the nucleosome, the length of DNA wrapped around the core, termed the wrapping efficiency. This value was obtained by subtracting the lengths of the DNA not wrapped around the nucleosome core from the total length of the DNA. As shown in Figure 2B, the wrapping efficiency for the H3_nuc_ sample is 146 ± 1.6 bp (SEM), which is in line with previous measurements on different DNA substrates, including the nucleosome-specific 601 motif ^30^.

**Figure 2.**
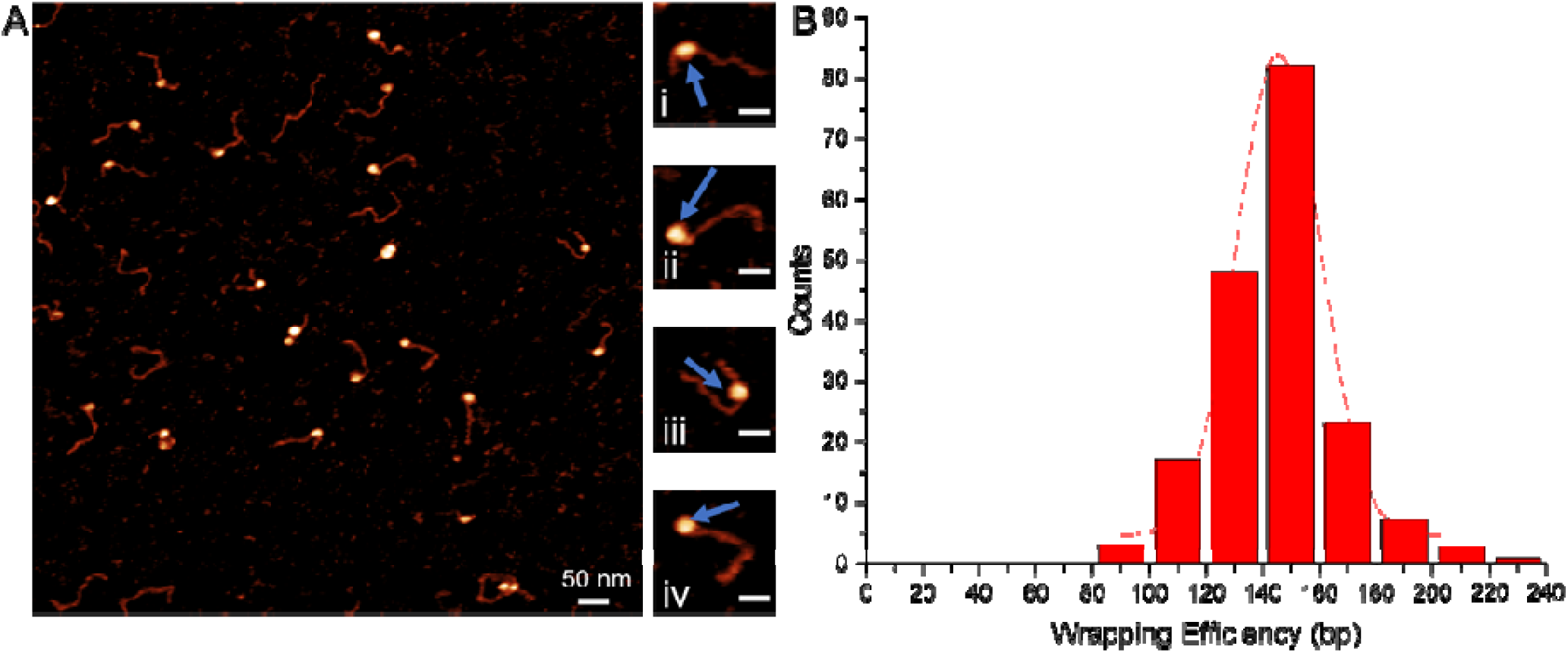
AFM image with zoomed-in snapshots of canonical H3_nuc_. AFM image (A) of the canonical H3_nuc_ assembled on the DNA construct. The large AFM image is a 1 x 1 μm scan size with a 50 nm scale bar. The snapshots to the right of the large AFM show the varying nucleosome binding locations. The snapshots are 100 x 100 nm scan area and 25 nm scale bars. Snapshots (ii and iv) show terminally bound nucleosomes, and (i and iii) are closer to the middle but are not terminally bound. The blue arrows indicate the location of a nucleosome. The wrapping efficiency of the nucleosomes on the DNA substrate was 146 ± 1.6 bp (SEM), as seen in the histogram (B).

Next, we added NF-κB_FL_ to the assembled H3_nuc_ sample in a 1:1 nucleosome:protein ratio, incubated the mixture for 10 min, and prepared the sample for AFM as in previous studies. AFM images are shown in Figure 3A, in which selected typical images of the complexes are shown to the right of the large AFM scan. Frame (i) shows just an H3_nuc_ (blue arrows), frames (ii and iv) show a terminally bound H3_nuc_ with an NF-κB_FL_ protein adjacent to it (orange arrows), and frame (iii) shows a terminally bound nucleosome with an NF-κB_FL_ on the opposite end of the DNA. The nucleosomes and NF-κB_FL_ can be visually differentiated based on their overall sizes, with NF-κB being significantly smaller than the nucleosome.

**Figure 3.**
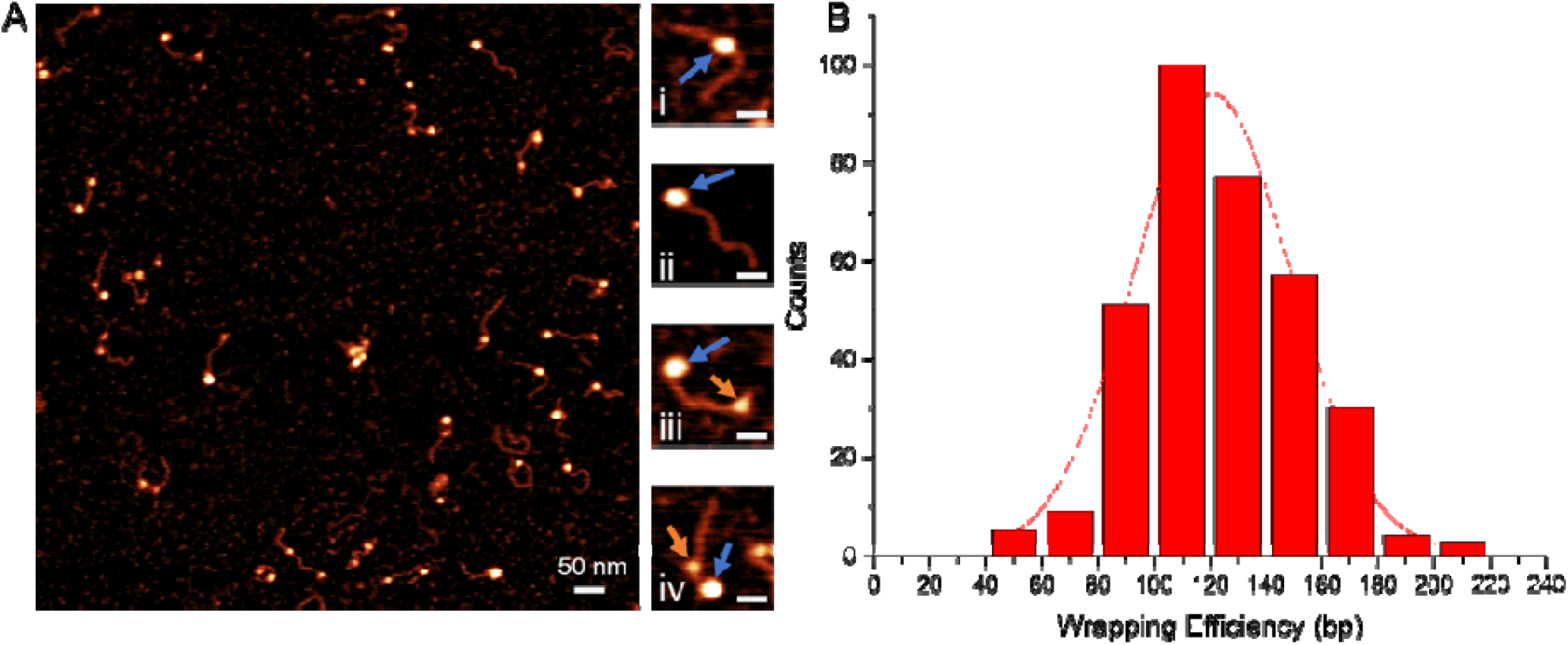
AFM image with zoomed-in snapshots of canonical H3_nuc_ with NF-κB_FL_. AFM images of H3_nuc_ assembled on the DNA construct with NF-κB_FL_ added at a 1 to 1 ratio(A). The large AFM image is a 1 x 1 μm scan size with a 50 nm scale bar. The snapshots to the right of the larger AFM image show varying situations. The snapshots are 100 x 100 nm scan area and 25 nm scale bars. In (i and ii), there is a nucleosome bound near the center of the DNA, with no NF-κB visible. In (iii), the nucleosome is bound to one side of the DNA, and the NF-κB i bound to the other side. In (iv), there is a terminally bound nucleosome with an NF-κB_FL_ bound adjacent. The orange arrows indicate NF-κB_FL_ bound to the DNA, and the blue arrows indicate the nucleosome. The wrapping efficiency was found to be decreased to 125 ± 2.2 bp (SEM), a seen in (B).

Next, we calculated the wrapping efficiency from these data as described above. The histogram from multiple measurements is shown in Figure 3B. The distribution was fit to a Gaussian distribution, yielding a mean value of the wrapping efficiency of 125 ± 2.2 bp (SEM). This number is considerably lower than the wrapping efficiency of the control sample, 146 ± 1.6 bp (SEM). This suggests that in the presence of NF-κB_FL,_ the nucleosomes are unraveled by some 21 ± 3.8 bp (SEM). The p-value between these two populations was 9.6 × 10^-10^, indicating a statistically significant difference between the control and the NF-κB_FL_ containing population.

We completed a parallel experiment that tested the effects of adding additional NF-κB to the nucleosome sample. In these experiments, we had a nucleosome: NF-κB ratio of 1 to 2 to check if increasing the concentration would affect the unwrapping effect of NF-κB. The AFM image can be seen in Figure 4A. In frame (i), a single nucleosome bound near the middle of the DNA can be seen. In frames (ii and iii), there is one H3_nuc_ and one NF-κB_FL_. In frame (iv), there is an H3_nuc_ and two NF-κB_FL_ bound to both sides of the nucleosome. The histogram of the wrapping results can be seen in Figure 4B. The unwrapping effect of the nucleosomes resulted in a wrapping efficiency of 125 ± 2.6 bp (SEM) regardless of the increased ratio of NF-κB. The yield of NF-κB bound to DNA flanks on the same strand as a nucleosome was calculated to be 43% and 85% for 1 to 1 and 1 to 2, respectively, which is in line with the use of higher concentration of the NF-κB.

**Figure 4.**
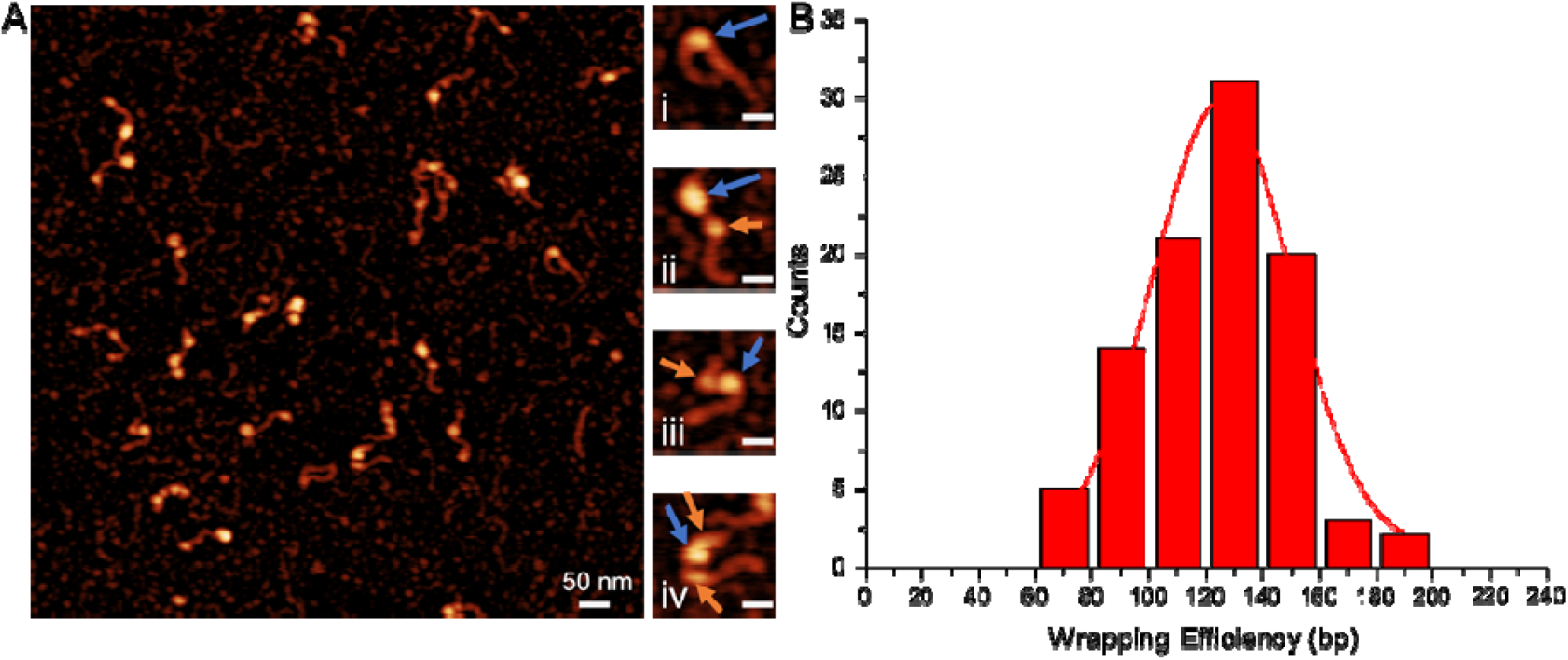
AFM image with zoomed-in snapshots of canonical H3_nuc_ with NF-κB_FL_. AFM images of H3_nuc_ assembled on the DNA construct with NF-κB_FL_ added at a 1 to 2 ratio (A). The large scan in (A) is 1 x 1 um, and the scale bar is 50 nm. The snapshots to the right of the larger AFM image show varying situations. The snapshots are 100 x 100 nm, and the scale bar is 25 nm. In (i), there is a nucleosome bound near the center of the DNA. In (ii and iii), the nucleosome is bound to the DNA, and the NF-κB is bound near the nucleosome. In (iv), the nucleosome has an NF-κB_FL_ bound to both sides of the nucleosome. The orange arrows indicate NF-κB_FL_ bound to the DNA, and the blue arrows indicate the nucleosome. The wrapping efficiency was found to be decreased to 125 ± 2.6 bp (SEM), as seen in (B).

Although NF-κB bound to the nucleosome itself cannot be visualized with AFM directly, its contribution to the particle size can be evaluated with AFM by the height or volume measurements. ^36^ The height of NF-κB_FL_ bound to DNA was measured, as well as the DNA height on each AFM image. The average DNA height was subtracted from the height measurements of NF-κB_FL_ bound to the DNA and was found to be 0.55 ± 0.02 nm (SEM) (n = 82). The height measurements for the set of 183 particles for the H3_nuc_ control produced the value 1.9 ± 0.02 nm (SEM). Similar measurements for the nucleosome particles (n = 157) in the presence of NF-κB led to the value 2.3 ± 0.04 nm (SEM), which is statistically significant from the control measurements. The p-value between these two populations was 5.4 x 10^-23^, indicating a statistically significant difference between the control and the NF-κB_FL_ containing population. These data are in Fig. S2A-C and summarized in Table 1.

**Table 1.** Results for experiments with both H3_nuc_ and CENP-A_nuc_ with and without the full length NF-κB for the 1:1 molar ratio.

|  | **H3 Nucleosomes** |  | **CENPA Nucleosomes** |  |
| --- | --- | --- | --- | --- |
| | Control | NF-κB<sub>FL</sub> | Control | NF-κB<sub>FL</sub> |
| Wrapping Mean (bp) | 146 ± 1.6 | 125 ± 2.2 | 130 ± 1.4 | 129 ± 2.2 |
| Height Mean (nm) | 1.9 ± 0.02 | 2.3 ± 0.04 | 2.2 ± 0.02 | 2.2 ± 0.04 |
| Volume Mean (nm<sup>3</sup>) | 355 ± 9 | 469 ± 9 | 351 ± 5 | 338 ± 8 |

We also completed a volume analysis of the control H3_nuc_ and the H3_nuc_ in the presence of NF-κB_FL_ and found that 355 ± 9.4 nm^3^ (SEM) and 469 ± 9.0 nm^3^ (SEM), respectively. A histogram distribution of these results can be seen in Supplementary Figures S3A, B. The p-value between these two populations was 1.8 x 10^-15^, indicating a statistically significant difference between the control and the NF-κB_FL_ containing population.

Similar studies were performed with truncated NF-κB_RHD_ protein. Images of the sample with snapshots can be seen in Supplementary Figure S4A, where the nucleosomes are indicated with a blue arrow, and the NF-κB_RHD_ are marked with orange arrows. The snapshots from the larger AFM image can be seen to the right, where frames (i and iii) show nucleosomes in different places on the DNA. In frames (ii and iv), a nucleosome is either terminally bound or close to the end of the DNA with an NF-κB_RHD_ protein bound on the DNA flank. The wrapping efficiency of the complex was decreased to 126 ± 1.5 bp (SEM) (Supplementary Fig. S4B). These data suggest that the C-terminal RelA TAD does not contribute to the unraveling property of NF-κB. The height measurements for the NF-κB_RHD_-bound nucleosomes was 1.9 ± 0.02 nm (SEM)(Supplementary Fig. S3C). The control for the subpopulation of H3_nuc_ with 130 bp wrapping efficiency resulted in a height of 1.67 ± 0.05 nm (SEM). This value is less than the height of complexes of NF-κB_RHD_ protein with nucleosome, suggesting that the truncated NF-κB_RHD_ is bound to the nucleosome.

Therefore, NF-κB leads to a substantial unraveling of H3_nuc_. The unraveling of nucleosome by NF-κB was reported in our recent publication ^30^, in which the nucleosome-specific 601 motif was used, but in this case, NF-κB unwrapped the 601 DNA to a lesser extent, only 135 ± 3 bp (SEM). Our data obtained on the physiologically-relevant DNA substrate indicates that the nucleosome unraveling is a property of NF-κB, but the effect quantitatively depends on the DNA sequence.

### NF-κB interaction with centromeric CENP-A_nuc_

Centromeric-specific CENP-A_nuc_ were assembled on the α-sat DNA substrate mentioned above (Fig. 1A) in a similar manner as the H3_nuc,_ with the exception that the CENP-A_nuc_ requires an extra step in the self-assembly process of mixing 2:1 molar concentrations of the dimeric H2A/H2B and tetrameric CENP-A/H4 (see the methods section for details). The AFM images of the nucleosomes are shown in Figure 5A, with a few snapshots selected to the right. The snapshots of the assembled CENP-A nucleosomes shown in frames (i and ii) show nucleosomes bound close to the DNA ends, and frames (iii and iv) show nucleosomes closer to the middle of the DNA sequence. The nucleosomes are indicated with blue arrows. The wrapping efficiency of the CENP-A_nuc_ was 130 ± 1.6 bp (SEM) (Fig. 5B), which is in line with our previous publications in which 601 motif DNA substrate along with non-specific DNA sequences were used. ^23,24,30^

**Figure 5.**
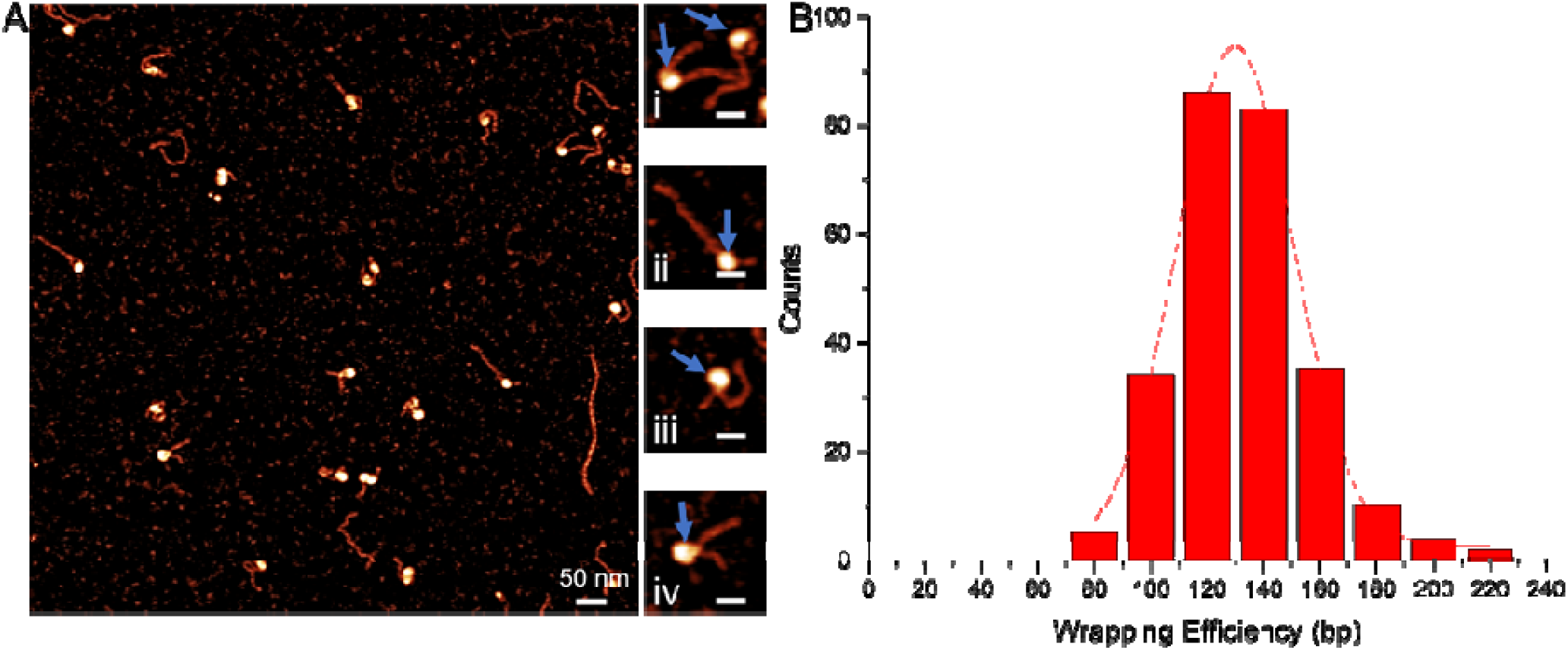
AFM image with zoomed-in snapshots of centromeric CENP-A_nuc_. AFM image of CENP-A_nuc_ assembled on the DNA construct is shown in (A). The large AFM image is 1 x 1 um, and the snapshots are 100 x 100 nm. The scale bars are 50 and 25 nm for the large AFM image and snapshots, respectively. The snapshots to the right of the large AFM image show typical nucleosomes assembled on the DNA construct. In (i), there are two nucleosomes, one close to the end and one more centrally bound. In (ii) and (iii), nucleosomes are close to the terminal end. In (iv), the nucleosome is closer to the middle of the DNA. The histogram to the right (B) represented the Gaussian distribution of the wrapping efficiency of the assembly, which was 130 ± 1.4 bp (SEM). The scale bar is 50 and 25 nm for the large image and snapshots, respectively.

Next, NF-κB_FL_ was added to CENP-A_nuc_ in a nucleosome-to-protein molar ratio of 1:1. The AFM results from the CENP-A_nuc_ and NF-κB_FL_ can be seen in Figure 6A, where selected complexes can be seen in the snapshots shown to the right of the large AFM image. In frame (i), the CENP-A_nuc_ is bound near the end of the DNA, and the NF-κB_FL_ is adjacent to the CENP- A_nuc_. In frame (ii), the CENP-A_nuc_ is bound to the middle, with the NF-κB_FL_ bound to the end of the DNA. In frame (iii), the CENP-A_nuc_ and the NF-κB_FL_ are bound to opposite ends of the DNA. The frame (iv) shows a CENP-A_nuc_ bound to the middle of the DNA, with no NF-κB_FL_ on the DNA flanks. The blue arrows indicate CENP-A_nuc_ and the orange arrows indicate the NF- κB_FL_ bound to DNA flanks. The wrapping efficiency of the nucleosomes at a 1 to 1 ratio was calculated as described above, and the histograms of multiple measurements can be seen in Figure 6B. The mean wrapping efficiency was 129 ± 1.6 bp (SEM). The wrapping efficiency of the nucleosomes at a 1 to 2 ratio is shown in Supplementary Figure S5A. In frame (i), a single CENP-A_nuc_ is bound to the DNA. In frames (ii and iii), there is a single CENP-A_nuc_ bound near the end of the DNA, with a single NF-κB_FL_ bound on the flank of the DNA. In frame (iv), there is an NF-κB_FL_ bound to both sides of the CENP-A_nuc_. The wrapping efficiency of the 1 to 2 experiments can be seen in Supplementary Figure S5B, which resulted in a wrapping efficiency of 131 ± 2.3 bp (SEM). Both the 1 to 1 and the 1 to 2 wrapping efficiencies were unchanged from the results found in the control sample, suggesting that in the presence of NF-κB, there is no unwrapping to the CENP-A_nuc_. The number of NF-κB on the DNA flanks with the CENP-A_nuc_ was 46% and 80% for molar ratios of 1:1 and 1:2, respectively. The p-value between the control and the 1 to 1 NF-κB_FL_ wrapping populations was 0.49, indicating no difference between the control and the NF-κB_FL_ populations. This increased binding of NF-κB to DNA indicates at least twice as many NF-κB seen bound to the DNA, which does not include the NF-κB that is potentially bound to the nucleosomes.

**Figure 6.**
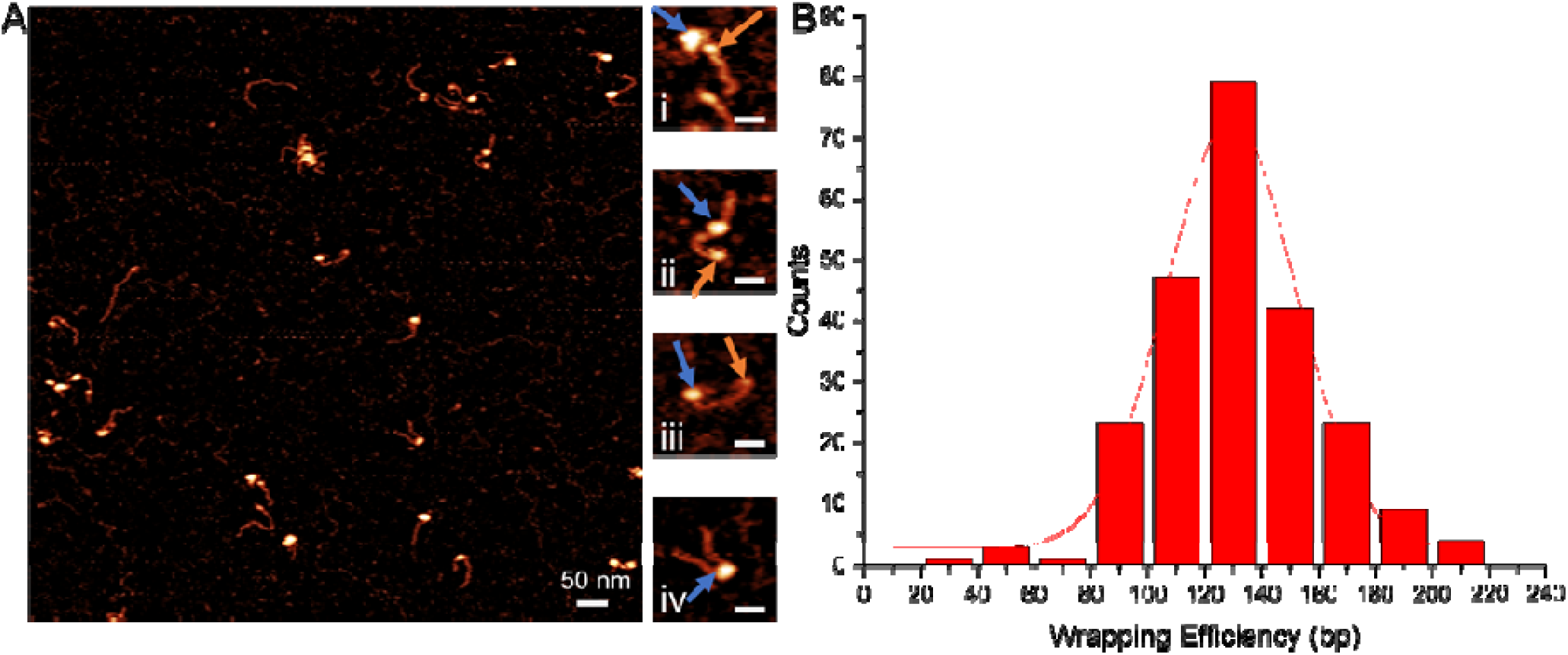
AFM image with zoomed-in snapshots of centromeric CENP-A_nuc_ with NF-κB_FL_. AFM image of CENP-A_nuc_ assembled on the DNA construct with 1 to 1 NF-κB_FL_ on the left (A) and histogram of wrapping efficiency on the right (B). The larger AFM image has a scan size of 1 x 1 um, and the snapshots are 100 x 100 nm and scale bars of 50 nm and 25 nm, respectively. The snapshots to the right of the large AFM image have NF-κB_FL_ added to the assembled nucleosomes and can be seen easily, represented by the orange arrows. The blue arrows represent the nucleosomes. The snapshots show nucleosomes binding near the terminal and can be seen in (i and iii), whereas centrally bound nucleosomes can be seen in (ii and iv). The CENP-A_nuc_ wrapping efficiency was 129 ± 1.6 bp (SEM).

Next, we looked for evidence of NF-κB_FL_ binding to the CENP-A_nuc_ through analysis of the nucleosome’s measured heights, which can be seen in Supplementary Figure S6A-C. The NF-κB_FL_ protein bound to the DNA, minus the height of the DNA, was measured and found to be 0.54 ± 0.03 nm (SEM). The CENP-A_nuc_ control nucleosomes had a height of 2.2 ± 0.02 nm (SEM), and with the addition of NF-κB_FL,_ the height was 2.2 ± 0.04 nm (SEM). The p-value between these two populations was 0.008, indicating little statistically significant difference between the control and the NF-κB_FL_ containing population. These results differ from the H3_nuc_, where an increase could be seen from the control upon the addition of NF-κB_FL_. With the CENP-A_nuc_, there was no increase in height with the addition of NF-κB_FL_, indicating no NF-κB_FL_ binding to the CENP-A_nuc_.

We also completed a volume analysis of the control H3_nuc_ and the H3_nuc_ in the presence of NF-κB_FL_ and found that 351 ± 8.2 nm^3^ (SEM) and 338 ± 4.9 nm^3^ (SEM), respectively. A histogram distribution of these results can be seen in Supplementary Figure S3C, D. The p-value between these two populations was 0.27, indicating little statistical difference between the control and the NF-κB_FL_ containing population.

Similar experiments were completed with the modified NF-κB_RHD_. Images and snapshots can be seen in Supplementary Figure S7A. The snapshots can be seen to the right of the large AFM image. In frames (i and ii), the CENP-A_nuc_ is bound to the DNA without any NF-κB_RHD_ bound to the flanks. In frames (iii and iv), the CENP-A_nuc_ are bound near the terminal end of the DNA with the NF-κB_RHD_ bound adjacent to the CENP-A_nuc_. The nucleosomes are indicated with a blue arrow, and the NF-κB_RHD_ are marked with orange arrows. The wrapping efficiency of CENP-A_nuc_ with NF-κB_RHD_ was 129 ± 1.6 bp (SEM)(Supplementary Fig. S7B), indicating no unwrapping. The height values of the nucleosomes were measured for the NF-κB_RHD_ containing samples and found to be 1.9 ± 0.02 nm (SEM)(Supplementary Fig. S7C) for CENP-A_nuc_. These results show that NF-κB does not unravel CENP-A_nuc_ nor bind to the nucleosome.

### Comparison of nucleosome positioning with NF-κB

We mapped the position of the nucleosome in these complexes, as shown in Figure 7A-C. Interestingly, the data show a lower population of nucleosomes at the DNA end when NF-κB i bound. This value was reduced to 16 % compared with 27 % for the control, which decreased even more with a 1 to 2 ratio decreasing to 9%. These results suggest that the NF-κB can cause displacement of the nucleosomes from the end of the DNA.

**Figure 7.**
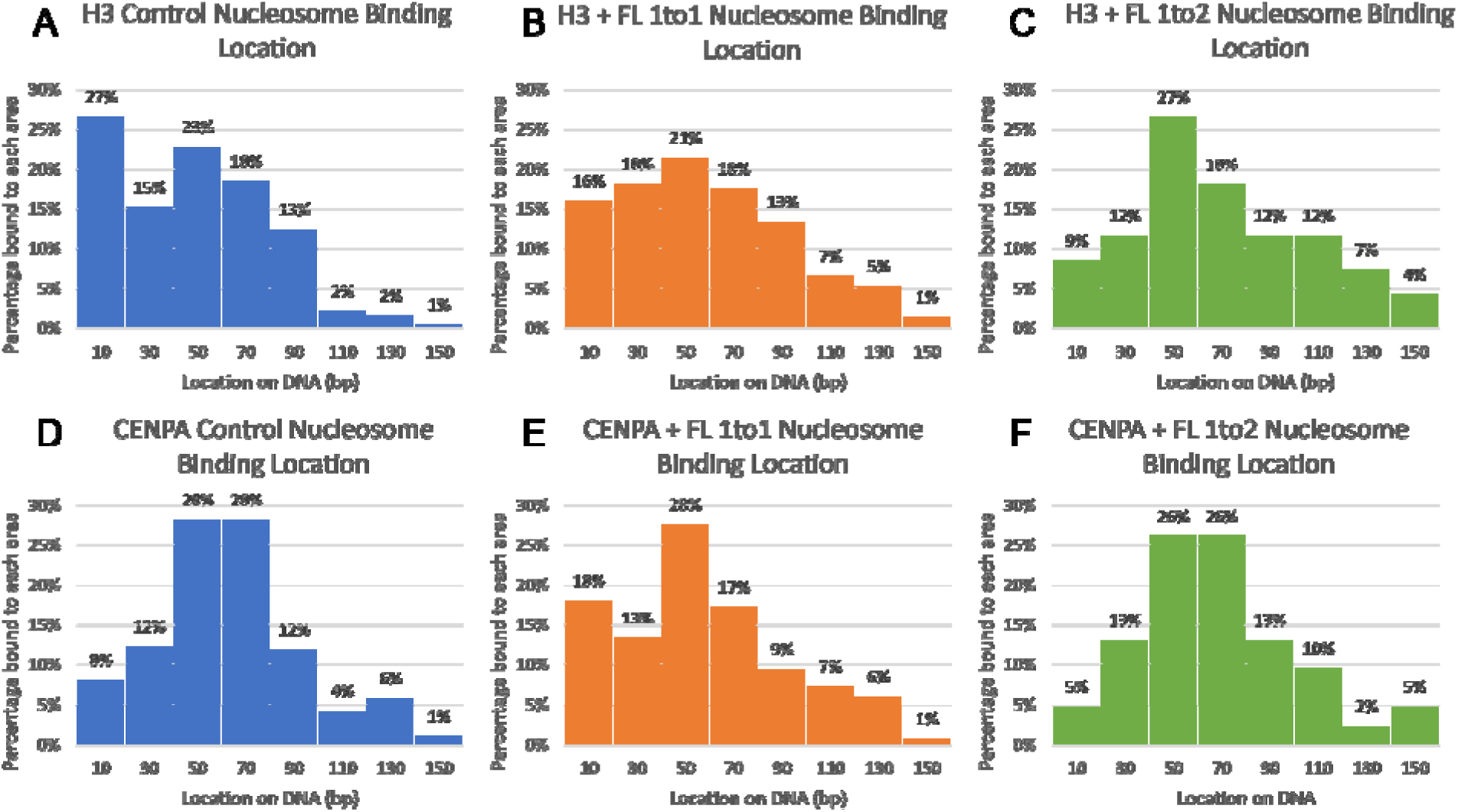
Nucleosome binding locations with varying concentrations of NF-κB_FL_. Canonical nucleosome binding locations along the DNA construct: H3_nuc_ Control (A), H3_nuc_ with 1 to 1 NF- κB_FL_ (B), and H3_nuc_ with 1 to 2 NF-κB_FL_ (C). The H3_nuc_ control had a terminal binding percentage of 27%, drastically decreasing to 16% in samples with NF-κB_FL_. The 1 to 2 NF-κB_F_ had an end binding of 9%, an even greater decrease in terminal binding than the 1 to 1 sample. Centromeric nucleosome binding locations along the DNA construct: CENP-A_nuc_ control (D), CENP-A_nuc_ with 1 to 1 NF-κB_FL_ (E), and CENP-A_nuc_ with 1 to 2 NF-κB_FL_ (F). The CENP-A_nuc_ control had an end binding percentage of 8%, which was increased to 18% in samples with NF-κB_FL,_ at a 1 to 2 ratio, the terminal bound CENP-A_nuc_ decreased to 5%.

The mapping results of the CENP-A_nuc_ control and in the presence of NF-κB_FL_ can be seen in Figure 7D-F. There was no consistent change in the mapping profile for the CENP-A nucleosomes in the presence of NF-κB.

The mapping data of H3_nuc_ with NF-κB_RHD_ added is shown in Supplementary Figure S8A, which is very close to the data obtained for the full-length NF-κB. Similarly, the CENP-A mapping results with the NF-κB_RHD_ were comparable to the effects of NF-κB_FL_, with 12% terminal binding compared to 18% (Supplementary Fig. S8B).

An analysis of the height compared to the position of the nucleosome showed no correlation between the two. A scatter plot of the comparison for H3_nuc_ can be seen in Supplementary Figure S9A-F. This indicates that nucleosome repositioning is not solely occurring when NF-κB is bound to the nucleosome itself but can also occur when NF-κB is bound at the flanking DNA.

We considered the possibility that the observed repositioning was actually a result of nucleosome removal. The results show that whether NF-κB was not present (control) or was present at a nucleosome:protein ratio of 1:1 or 1:2, the yield of H3_nuc_ were 62%, 62%, and 63%, respectively. These results indicate that the NF-κB is not removing the nucleosomes from the DNA but rather causes translocation of the nucleosomes away from the DNA ends. The yield analysis was also completed for the CENP-A_nuc_; the results were 70%, 67%, and 71% for control, 1:1, and 1:2, respectively.

## Discussion

Our major findings are summarized in Figure 8 (see also Table 1). According to the graph in Figure 8, NF-κB unravels canonical H3 nucleosomes, removing more than 20 bp DNA out of 147 bp total DNA wrapped around the nucleosome core. The unraveling of nucleosomes by NF- κB was reported in our recent publication ^30^. However, nucleosomes were assembled on the highly specific 601 sequence in that publication, and a lower unwrapping effect was observed. Elevated stability of the nucleosome assembled by the specific 601 DNA sequence can explain this effect ^37^. Still, we need to consider the difference in the interaction of NF-κB with both DNA templates. The 601 motif contained only one κB binding site for NF-κB ^30^. However, analysis of the NF-κB binding data did not reveal a preference for NF-κB binding at that site ^30^. In contrast, the LJ-sat DNA substrate used in this work reveals a specific binding pattern of NF-κB (Fig. 1). Importantly, the protein showed higher affinity for the DNA ends as well as for a cluster of half κB sites near the middle of the sequence. The affinity of NF-κB to the DNA ends can explain the decrease in the population of the end-bound nucleosomes in the presence of NF-κB by 1.5 times (Fig. 7A-C). These observations suggest that binding of NF-κB to specific sites on DNA weakens nucleosome interactions, resulting in their displacement and/or dissociation. The findings in this paper on the similarity of nucleosome assembly on alpha-satellite and plasmid DNA sequences align with our previous findings ^23^. Here, we found the elevated affinity of nucleosomes to the DNA ends, which is greater than the affinity to the alpha-satellite sequence, which could be related to other properties of the nucleosomes, such as their dynamics.

**Figure 8.**
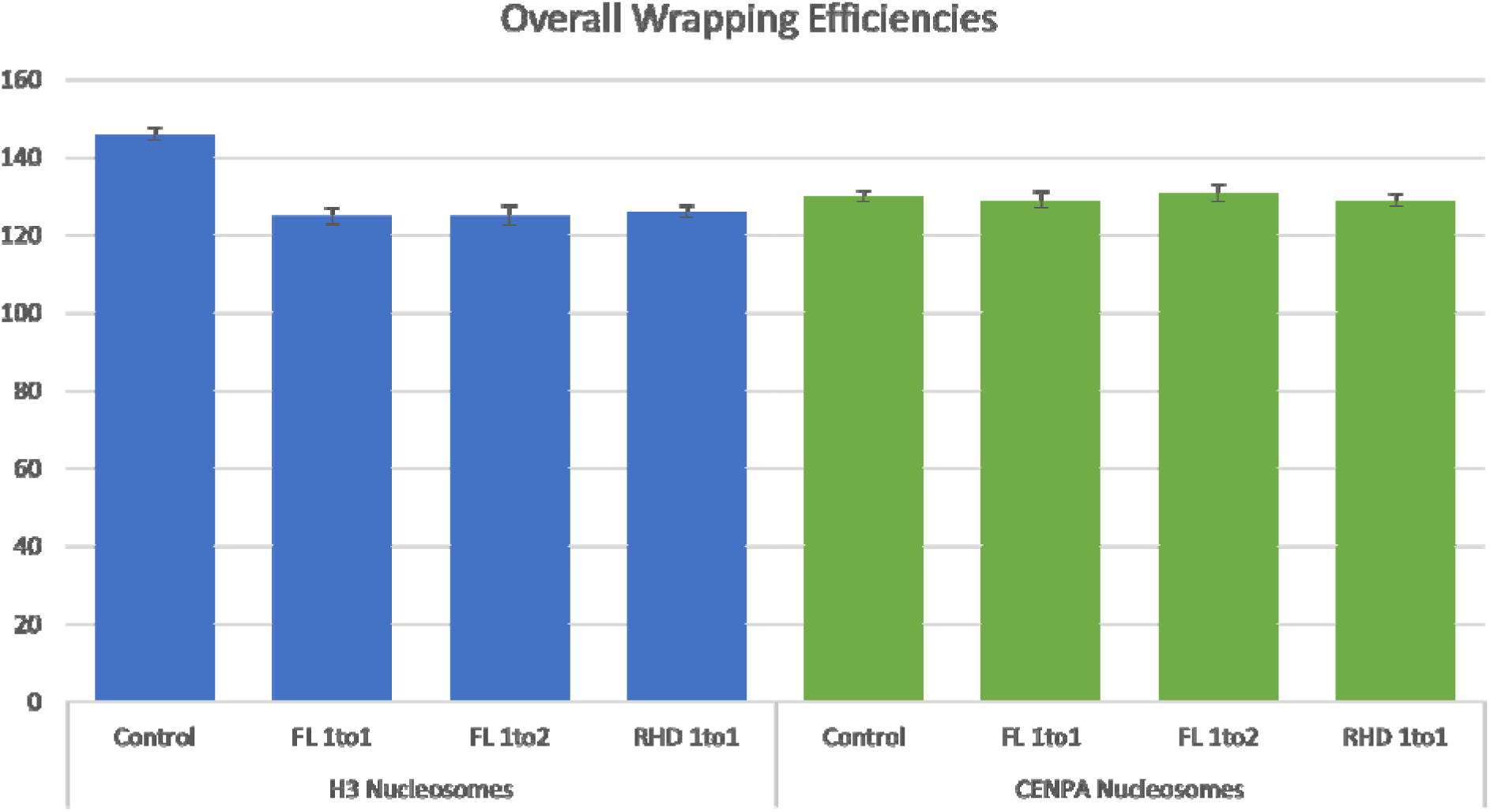
Effect of NF-κB on nucleosome wrapping efficiency. The H3_nuc_ control had the highest wrapping efficiency at 146 ± 1.6 (SEM) bp, whereas the H3_nuc_ with NF-κB_FL_ 1 to 1, NF- κB_FL_ 1 to 2, and NF-κB_RHD_ 1 to 1 had a lower wrapping at 125 ± 2.2 (SEM) bp, 125 ± 2.6 (SEM) bp, and 126 ± (SEM) 1.5 bp, respectively. The CENP-A_nuc_ results were very different from the H3_nuc_ with the NF-κB not affecting the wrapping efficiency: the CENP-A_nuc_ control (130 ± 1.3 [SEM] bp), with 1 to 1 NF-κB_FL_ (129 ± 2.2 [SEM] bp), with 1 to 2 NF-κB_FL_(131 ± 2.3 [SEM] bp), and with 1 to 1 NF-κB_RHD_ (129 ± 1.6 [SEM] bp). FL and RHD stand for the full-length and truncated variant NF-κB, respectively. The blue bars represent the H3_nuc_ wrapping, the green bars represent the CENP-A_nuc_ wrapping, and the error bars show the SEM.

Similar results of unraveling nucleosomes by NF-κB were obtained for the truncated variant NF-κB_RHD_ (Fig. S2), suggesting that the RelA C-terminal TAD does not define the unwrapping property of NF-κB. Given that the full-length NF-κB and its truncated variant NF-κB_RHD_ have similar DNA binding patterns (Fig. S1), we hypothesize that the DNA binding affinity of NF-κB is the factor defining the nucleosome unraveling property of NF-κB. We hypothesize that NF-κB binds to transiently dissociated DNA segments formed during the breathing of the nucleosome, stabilizing such an open state of the nucleosome and shifting the location of the nucleosome, explaining the repositioning of the population of the end-bound nucleosomes in the presence of NF-κB.

To look at the effect of the wrapping vs. the height of the nucleosomes both with and without NF-κB, we plotted them in Supplementary Figures S10 and S11. The trend can be seen that typically, a lower height is indicative of lower wrapping. The interaction of NF-κB with CENP-A nucleosomes is entirely different. As seen in Figures 3, 4, 6, and Supplementary Figure S4, there is no change in the nucleosome wrapping, suggesting that regardless of the same affinity of NF-κB to DNA, the protein cannot unravel the CENP-A nucleosome. A broader wrapping efficiency can be seen in the populations with NF-κB. This effect is likely from a small subpopulation of nucleosomes that remain canonically wrapped and have not had the NF-κB unwrapping effect. Therefore, we have two populations: the first, and larger population, is unwrapped due to NF-κB, and the second is the canonical wrapping. The combination of both populations has the effect of a broader histogram for the wrapping efficiency. Mixing of NF-κB with nucleosomes in the 1:1 ratio sometimes results in the binding of two proteins to one nucleosome; this is statistically what would be expected. Our experimental data clearly demonstrates this with complexes of the DNA substrate with NF-κB only. There were complexes with two or even three NF-κB bound to a single DNA and others with no NF-κB bound (Fig. 1), indicating that there can be 1 NF-κB bound to the DNA and one bound to the nucleosome, while other complexes have no NF-κB bound. In Figures 3, 4, and 6, we intentionally showed AFM images with NF-κB and nucleosomes visible as an internal control for NF-κB presence. The volume and height measurements carried out on hundreds of observed complexes show that NF-κB also binds to nucleosomes.

If breathing of nucleosomes is the pathway by which NF-κB unwraps the nucleosome, the NF-κB will bind to the transiently dissociated DNA segments. In that case, these data indicate that CENP-A nucleosomes are more stable than canonical H3 nucleosomes. According to the graph in Supplementary Figures S6B, C, there are no changes in the CENP-A nucleosome, suggesting that NF-κB does not bind the CENP-A nucleosome, which can be explained by the elevated stability of CENP-A nucleosomes compared with canonical ones ^24^. The ability of CENP-A nucleosomes to resist the binding of the high affinity of NF-κB to DNA can be a factor contributing to findings that in vitro CENP-A chromatin was predominantly nonpermissive for transcription compared to H3 chromatin ^38^. This finding is in line with previous results, which showed a 200-300 fold lower transcriptional activity, including that of NF-κB, of centromeric chromatin with euchromatin ^27^.

## ASSOCIATED CONTENT

### Supporting Information

Additional experimental AFM results and analysis, as well as height and volume analysis for both H3_nuc_ and CENP-A_nuc._ Comparative analysis of nucleosome height vs position on the DNA and height vs wrapping of nucleosomes. (DOC)

## AUTHOR INFORMATION

### Author Contributions

Shaun Filliaux designed research, performed research, analyzed data and wrote the paper. Chloe Bertelsen performed research and analyzed data. Hannah Baughman contributed protein and wrote the paper. Elizabeth Komives designed research, analyzed data and wrote the paper. Yuri Lyubchenko designed research, analyzed data and wrote the paper.

## ACKNOWLEDGMENT

We thank L. Shlyakhtenko (University of Nebraska Medical Center) for useful insights and all of the Y.L.L lab members for fruitful discussions of the data.

## ABBREVIATIONS

AFM: Atomic Force Microscopy
CENP-A_nuc_: CENP-A containing nucleosomes
H3_nuc_: H3 containing nucleosomes
NF-κB: Nuclear factor kappa-light-chain-enhancer of activated B cells
NF-κB_FL_: full length NF-κB
NF-κB_RHD_: NF-κB with the RHD removed
SEM: standard error of the mean
TAD: transcription activation domain;

