## supplemental data for "The Interaction of NF-κB Transcription Factor with Centromeric Chromatin"

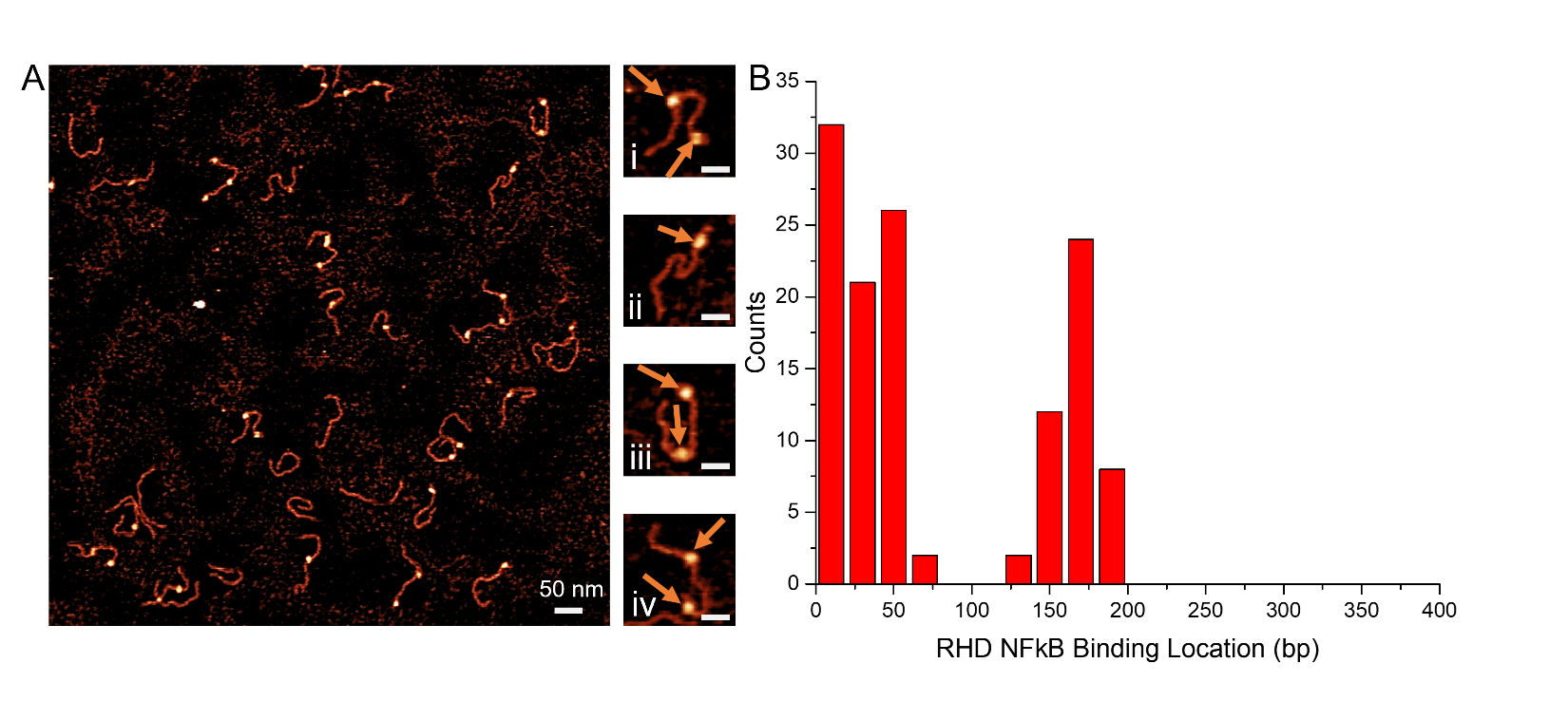
**Supplementary Figures**

**Supplementary Figure S1 – AFM image with zoomed-in snapshots of NF-κB_RHD_ results.** AFM images of NF-κB_RHD_ (A) on the DNA substrate. The orange arrows indicate an NF-κB_RHD_ bound to the DNA. The snapshots to the right of the large AFM image show various binding locations of individual NF-κB_RHD_. In the snapshots (i, iii, and iv), two nucleosomes are bound to the DNA, with one near the end of the DNA and the other near the middle. In snapshot (ii), a single protein is bound near the terminal end of the DNA. The large AFM image is a 1 μm x 1 μm scan size with a 50 nm scale bar. The snapshots are 100 x 100 nm scan area and 25 nm scale bars. Analysis of AFM images yielded the results of the NF-κB_RHD_ binding locations to have a preferential terminal binding when a single NF-κB is bound to the DNA (B) and a secondary peak near 170 bp.

**Supplementary**
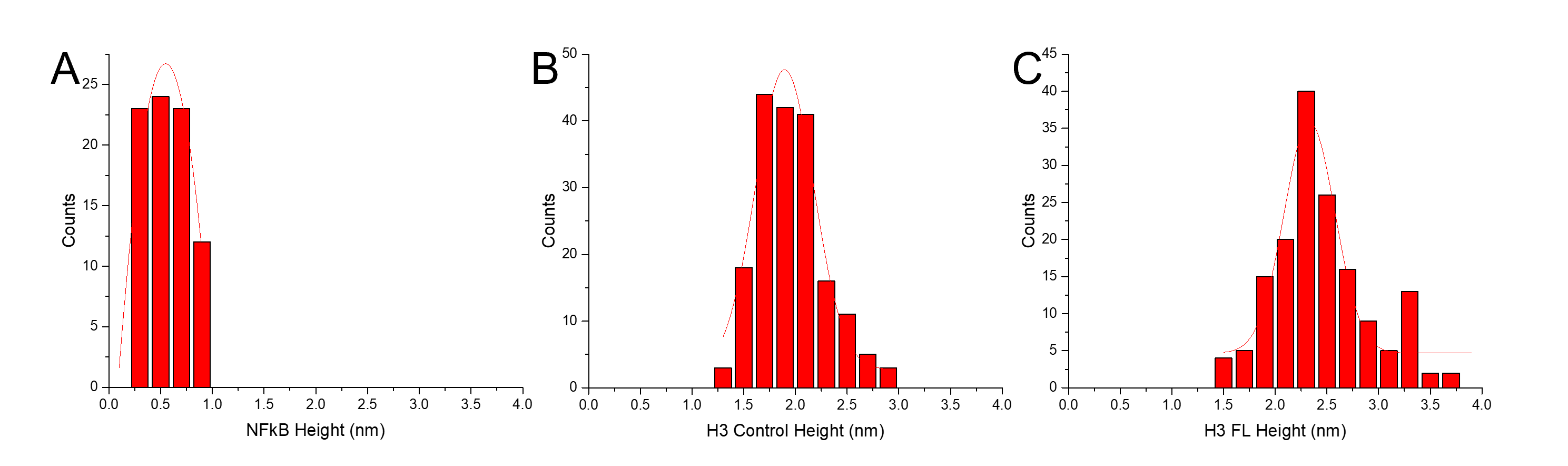
**Figure S2**. **Height values for H3_nuc,_ NF-κB_FL_ and its complexes of with H3_nuc._** The height results for the NF-κB_FL_ protein only (A), H3_nuc_ control (B), and H3_nuc_ in the presence of NF-κB_FL_ (C). Histograms were approximated with Gaussian distributions (red curves). The mean values for NF-κB_FL_ protein only, H3_nuc_ control, and H3_nuc_ in the presence of NF-κB_FL_ were 0.55 ± .02 nm, 1.9 ± 0.02, 2.3 ± 0.04 nm (SEM), respectively.

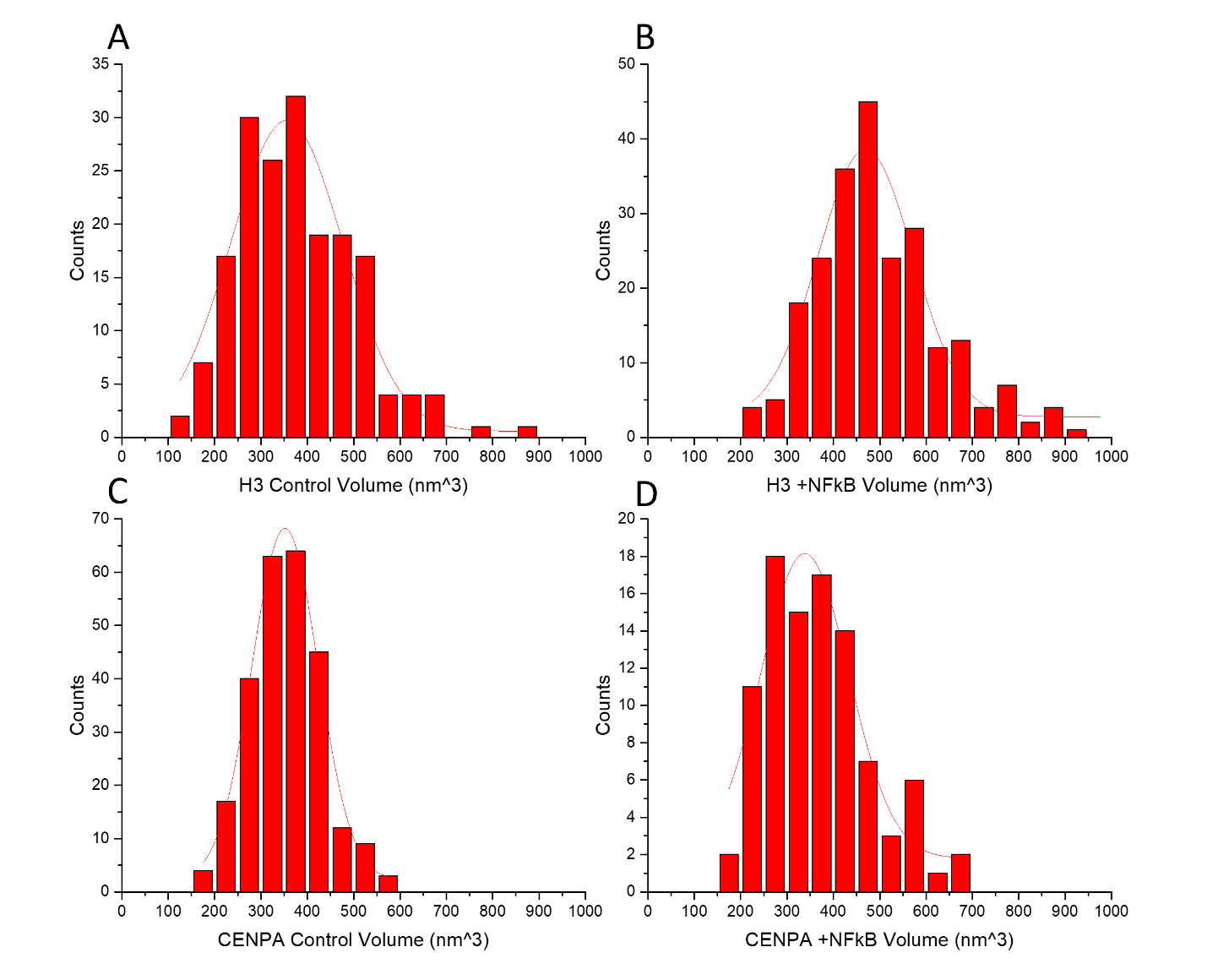
**Supplementary Figure S3 – Volume analysis of H3 and CENP-A nucleosomes.** The histograms show the results of the volume analysis of H3_nuc_ control (A), H3_nuc_ + NF-κB_FL_ (B), CENP-A_nuc_ control (C), and CENP-A_nuc_ + NF-κB_FL_ (D)**.** Histograms were approximated with Gaussian distributions (red curves).The average volume of the H3 Control, H3 + NF-κB_FL_, CENP-A control, and CENP-A + NF-κB_FL_ was 355 ± 9.4 nm^3^, 469 ± 9.0 nm^3^, 351 ± 4.9 nm^3^, and 338 ± 8.2 nm^3^ (SEM), respectively.

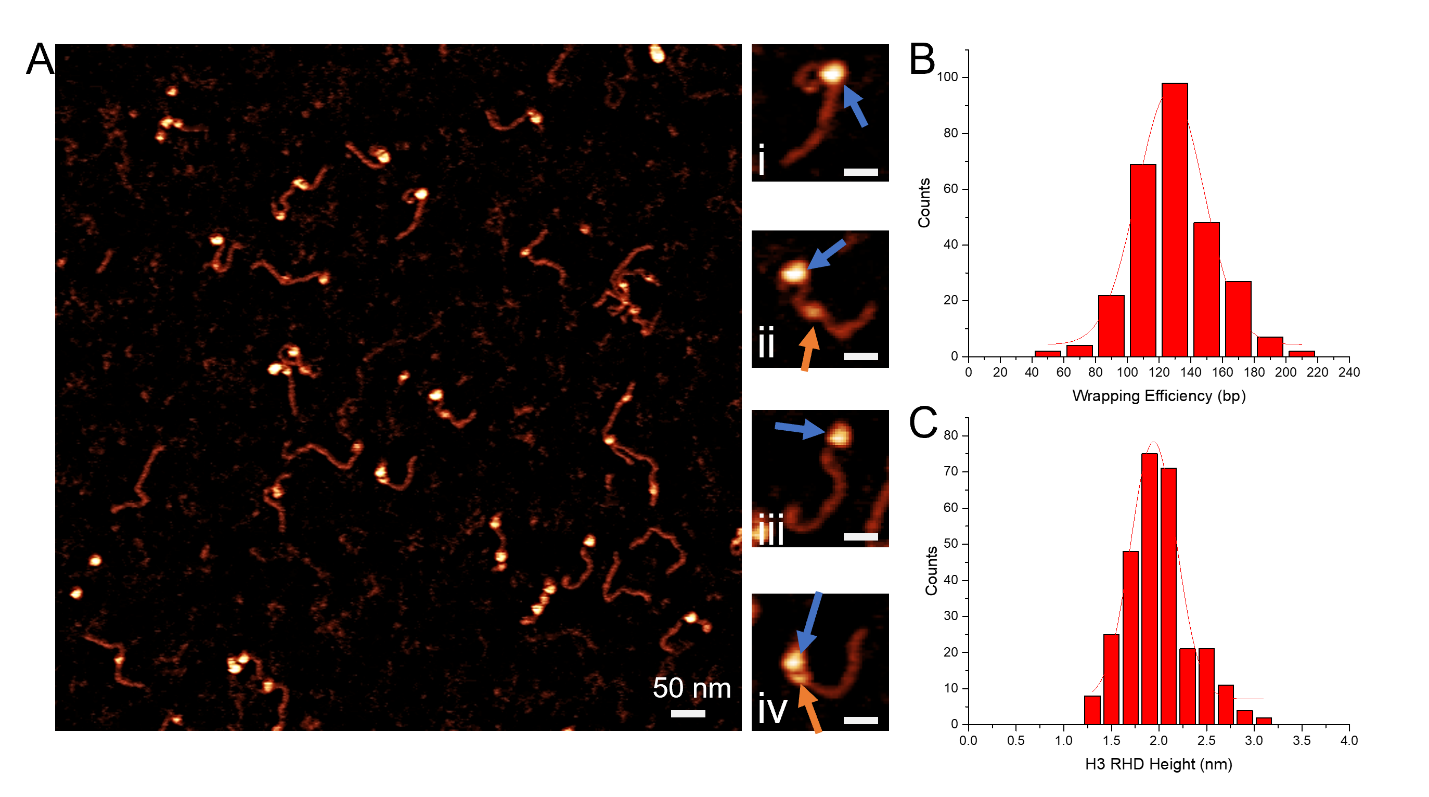
**Supplementary Figure S4 – AFM image with zoomed-in snapshots of canonical H3_nuc_ with NF-κB_RHD_.** AFM image of H3_nuc_ assembled on the DNA construct with added NF-κB_RHD_(A). Snapshots of the image scanned can be seen to the right of the large AFM image, with images (i and iii) showing a nucleosome bound to the DNA and in (ii and iv) a nucleosome bound with an NF-κB_RHD_ bound to the flank of the DNA. The large scan in (A) is 1 μm x 1 μm, and the scale bar is 50 nm. The snapshots are 100 x 100 nm, and the scale bar is 25 nm. The orange arrows indicate NF-κB_RHD_ bound to the DNA, and the blue arrows indicate the nucleosome. The wrapping efficiency was found to be decreased to 126 ± 1.5 bp (SEM), as seen in (B). Histograms were approximated with Gaussian distributions (red curves).The histograms shows the nucleosome height for H3_nuc_ with NF-κB_RHD_ (C). The mean height for H3_nuc_ with NF-κB_RHD_ was 1.9 ± 0.02 nm (SEM).

**Supplementary**
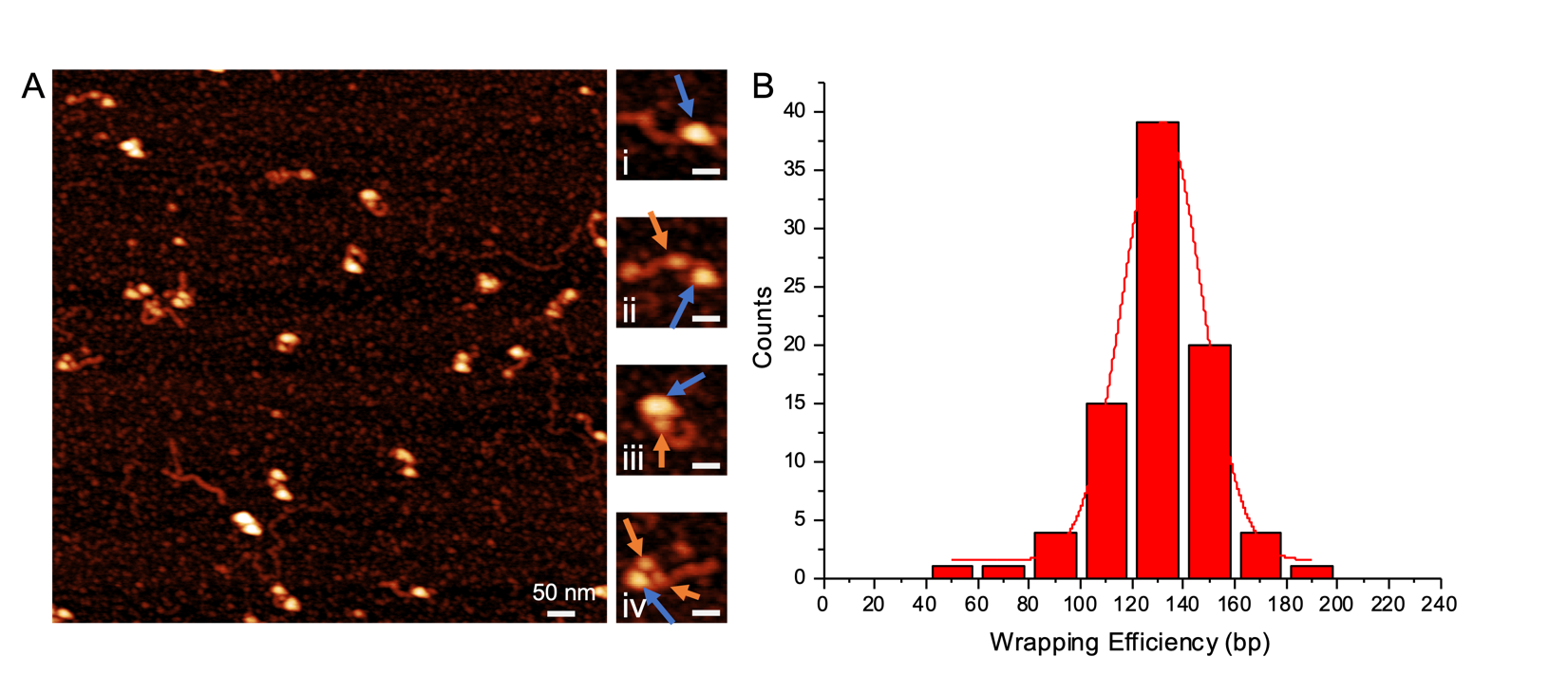
**Figure S5 -** **AFM image with zoomed-in snapshots of centromeric CENP-A_nuc_ with increased NF-κB_FL_ ratio**_._ AFM image of CENP-A_nuc_ assembled on the DNA construct with NF-κB_FL_ added at a 1:2 ratio with AFM images on the left (A) and histogram of wrapping efficiency on the right (B). Histograms were approximated with Gaussian distributions (red curves).The snapshots to the right of the larger AFM image show varying situations, NF-κB_FL_ represented by orange arrows. The blue arrows represent the nucleosomes—the snapshots to the right show various binding locations of the nucleosomes and NF-κB_FL_. In (i), there is a nucleosome bound near the end of the DNA. In (ii and iii), the nucleosome is bound to the DNA, and the NF-κB is bound near the nucleosome. In (iv), the nucleosome has an NF-κB_FL_ bound to both sides of the nucleosome. The large AFM image has a scan size of 1 μm x 1 μm, and the snapshots are 100 x 100 nm and a scale bar of 50 nm and 25 nm, respectively. The wrapping efficiency was 131 ± 2.3 bp (SEM).

**Supplementary**
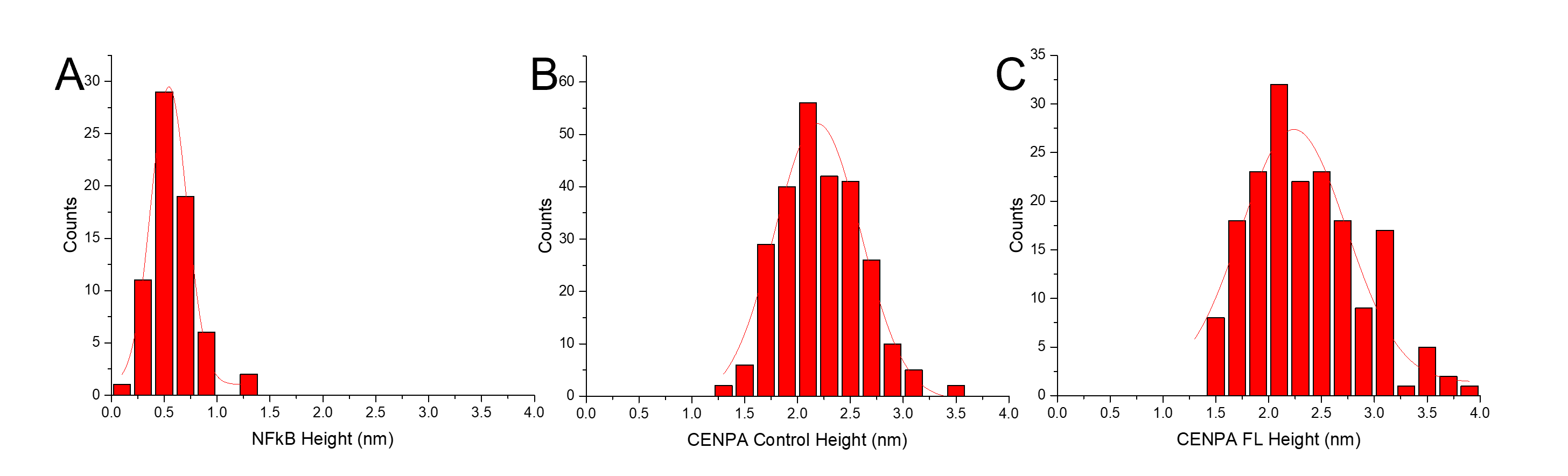
**Figure S6** – **Height results of NF-κB_FL_, CENP-A_nuc_, and CENP-A_nuc_ + NF-κB_FL._** The height results for NF-κB_FL_ protein only (A), CENP-A_nuc_ control (B), and CENP-A_nuc_ in the presence of NF-κB_FL_(C). The mean height of NF-κB_FL_ protein only, CENP-A_nuc_ control, and NF-κB_FL_ were 0.54 ± 0.03, 2.2 ± 0.02, and 2.2 ± 0.04 nm (SEM), respectively. Histograms were approximated with Gaussian distributions (red curves).

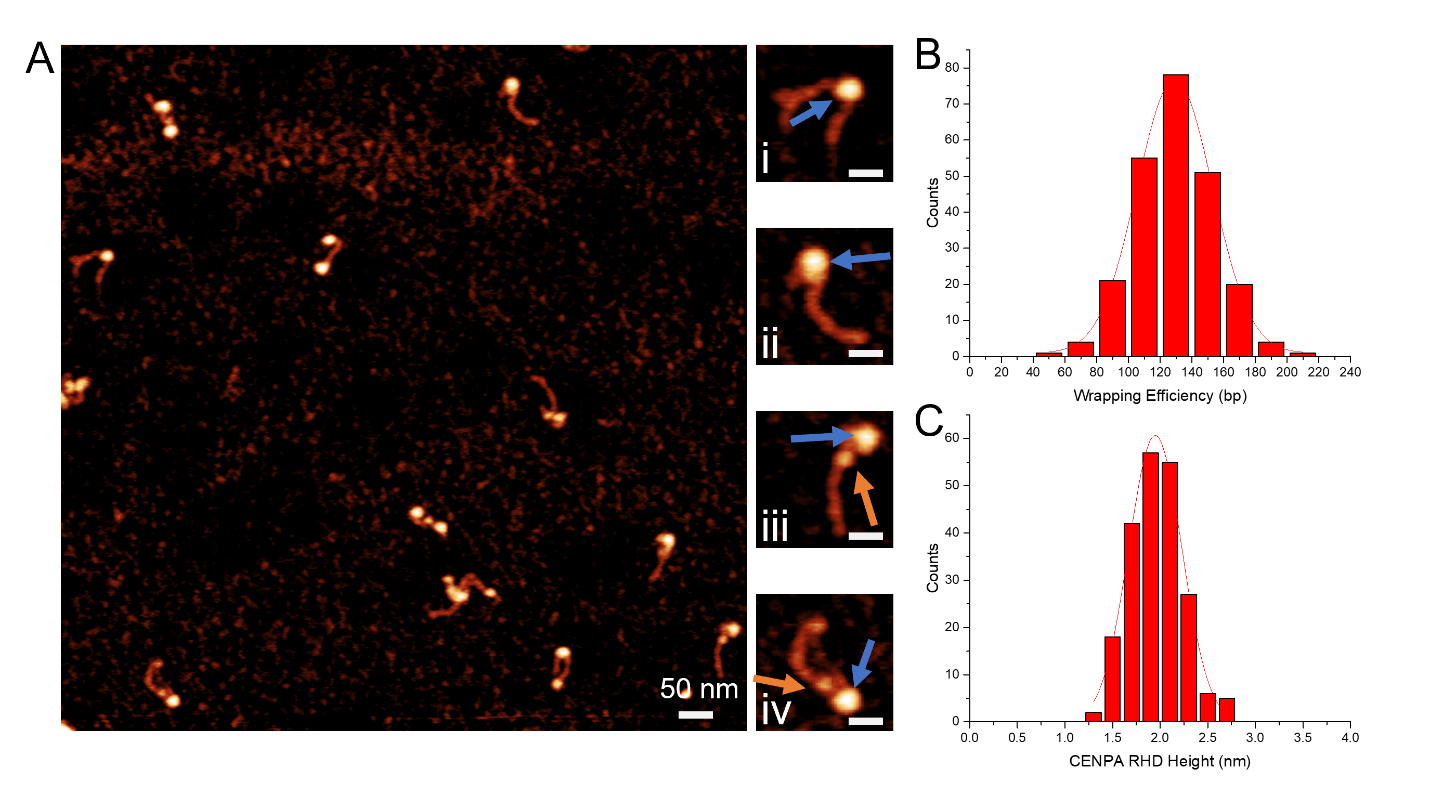
**Supplementary Figure S7 –** **AFM image with zoomed-in snapshots of centromeric CENP-Anuc with NF-κB_RHD_**. AFM image of CENP-A_nuc_ assembled on the DNA on the left (A) and histogram of wrapping efficiency in the upper right (B). NF-κB_RHD_ was added to the assembled nucleosomes and can be seen easily in the snapshots, represented by the orange arrows. In snapshots (i and ii), the nucleosome is bound to the DNA, with (i) being more centrally bound and (ii) being closer to the terminal end. In (iii and iv), terminal or nearly terminal bound nucleosomes have an adjacent NF-κB_RHD_ bound. The blue arrows represent the nucleosomes. The large AFM image has a scan size of 1 μm x 1 μm, and the snapshots have a size of 100 x 100 nm and a scale bar of 25 nm. The wrapping efficiency was 129 ± 1.6 bp (SEM)—the histogram showing the nucleosome height for CENP-A_nuc_ with NF-κB_RHD_ can be seen in (C). Histograms were approximated with Gaussian distributions (red curves). The mean height for CENP-A_nuc_ with NF-κB_RHD_ was 1.9 ± 0.02 nm (SEM).

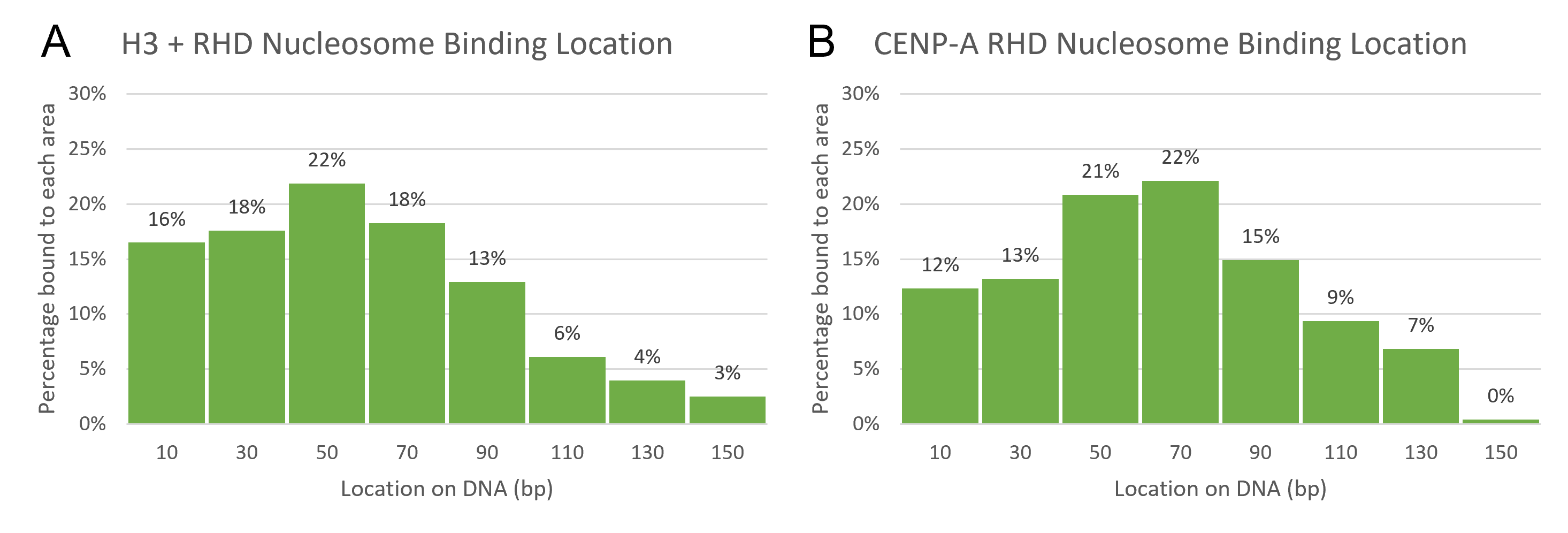

**Supplementary Figure S8** – **Nucleosome positioning at 1 to 1 nucleosome:protein of NF-κB_RHD_.** The results of the nucleosome positioning in the presence of NF-κB_RHD_ were similar to those achieved in the presence of NF-κB_FL_. The H3_nuc_ (A) decreased the end binding of the nucleosomes, and the CENP-A_nuc_ (B) had a slight increase in end binding.

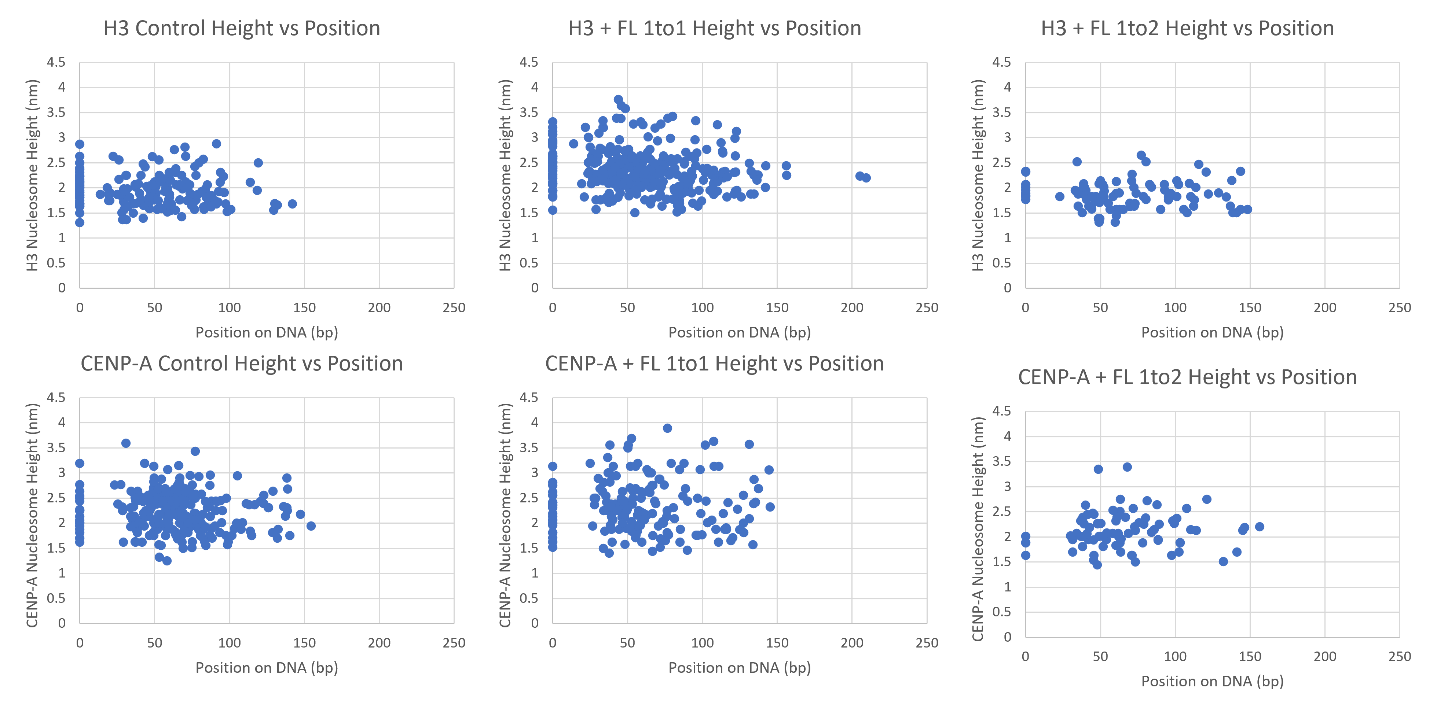

**Supplementary Figure S9 –** **Analysis of the nucleosome’s height as a comparison to the position on the DNA.** There was no correlation to the nucleosome height when compared to the position on the DNA; this includes the end binding, which can be seen at the 0 on the X-axis.

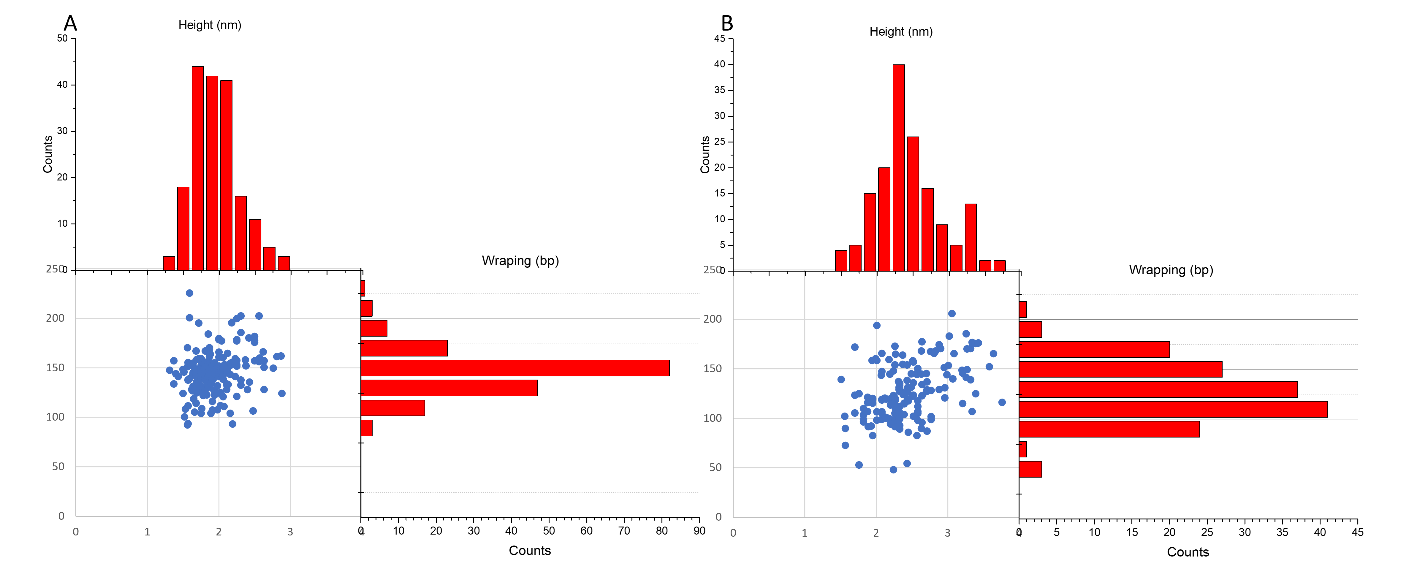

**Supplementary Figure S10 –** **Height vs Wrapping analysis for H3_nuc_.** Overall populational analysis of the height and wrapping efficiency of the H3_nuc_ samples. In frame A, the H3_nuc_ control sample had a mean wrapping of 146 ± 1.6 bp (SEM) and a mean height of 1.9 ± 0.02 nm (SEM). In frame B, the H3_nuc_ with NF-κB_FL_ had a mean wrapping of 125 ± 2.2 bp (SEM) and a mean height of 2.3 ± 0.04 nm (SEM). The X-axis is the height of each population, and the Y-axis is the wrapping efficiency of the nucleosomes.

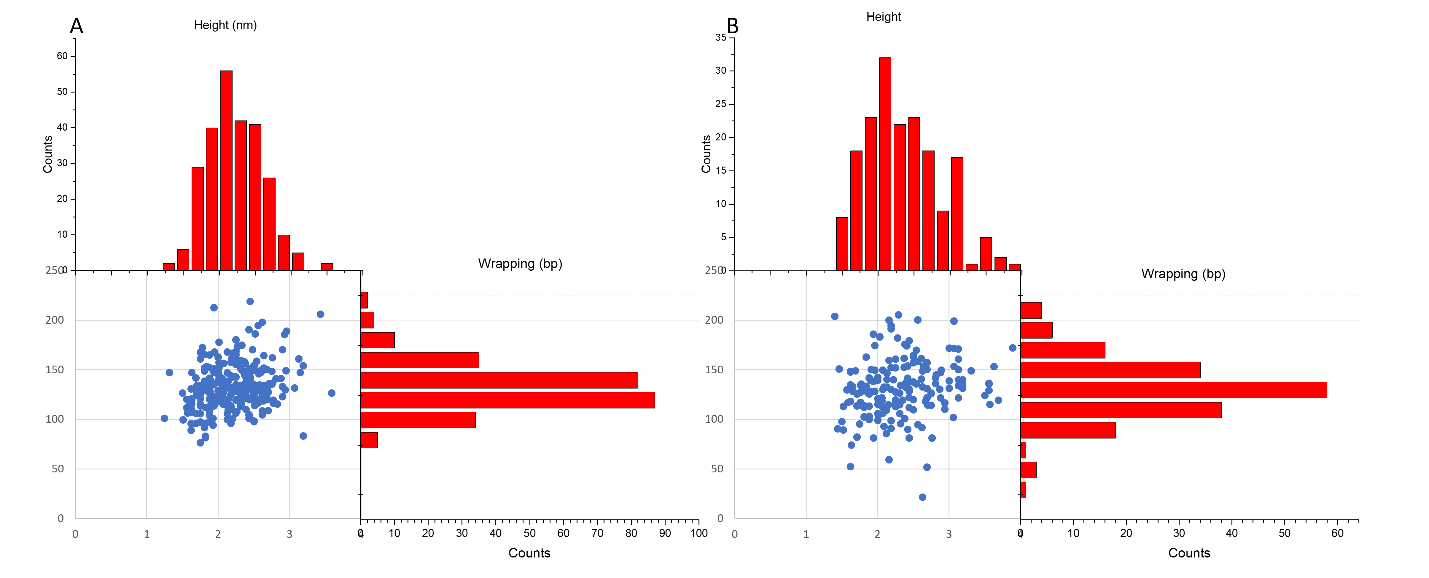
**Supplementary Figure S11 -** **Height vs Wrapping analysis for CENP-A_nuc_.** Overall populational analysis of the height and wrapping efficiency of the CENP-A_nuc_ samples. In frame A, the CENP-A_nuc_ control sample had a mean wrapping of 130 ± 1.6 bp (SEM) and a mean height of 2.2 ± 0.02 nm (SEM). In frame B, the CENP-A_nuc_ with NF-κB_FL_ had a mean wrapping of 129 ± 1.6 bp (SEM) and a mean height of 2.2 ± 0.04 nm (SEM). The X-axis is the height of each population, and the Y-axis is the wrapping efficiency of the nucleosomes.
